# Human Elp3/Kat9 is a mitochondrial tRNA modifying enzyme

**DOI:** 10.1101/2022.03.24.485634

**Authors:** Rachid Boutoual, Hyunsun Jo, Indra Heckenbach, Ritesh Tiwari, Herbert Kasler, Chad A. Lerner, Samah Shah, Birgit Schilling, Vincenzo Calvanese, Matthew J. Rardin, Morten Scheibye-Knudsen, Eric Verdin

## Abstract

Post-translational modifications, such as lysine acetylation, regulate the activity of diverse proteins across many cellular compartments. Protein deacetylation in mitochondria is catalyzed by the enzymatic activity of the NAD+-dependent deacetylase sirtuin 3 (SIRT3), however it remains unclear whether corresponding mitochondrial acetyltransferases exist. We used a bioinformatics approach to search for mitochondrial proteins with an acetyltransferase catalytic domain, and identified a novel splice variant of ELP3 (mt-ELP3) of the elongator complex, which localizes to the mitochondrial matrix in mammalian cells. Unexpectedly, mt-ELP3 does not mediate mitochondrial protein acetylation but instead induces a post-transcriptional modification of mitochondrial-transfer RNAs (mt-tRNAs). Overexpression of mt-ELP3 leads to the protection of mt-tRNAs against the tRNA-specific RNase angiogenin, increases mitochondrial translation, and furthermore increases expression of OXPHOS complexes. This study thus identifies mt-ELP3 as a non-canonical mt-tRNA modifying enzyme.

## Introduction

Lysine acetylation is a prominent regulator of multiple proteins that reside in different cellular compartments, including mitochondrial metabolic enzymes (Wagner & Payne, 2011). This reversible posttranslational modification is controlled by competing lysine acetyltransferases (KATs) and deacetylases (KDACs) that catalyze the addition and removal, respectively, of an acetyl group from a lysine residue (Carrico et al., 2018). We previously showed that SIRT3 is a mitochondrial NAD^+^-dependent protein deacetylase that controls energy metabolism (Björn Schwer et al., 2002). These findings motivated us to look for any corresponding mitochondrial KAT (s). Since the mitochondrial matrix has a higher acetyl-CoA concentration and a higher pH than other cellular compartments, mitochondrial acetylation has been proposed to occur via a non- enzymatic mechanism (Wagner & Payne, 2011). However, this mechanism does not exclude the existence of a mitochondrial KAT. Indeed, GCN5L1 (general control of amino acid synthesis 5)- like 1 and ACAT1 (acetyl-CoA acetyltransferase 1) are plausible mitochondrial KATs (Fan et al., 2014) (Scott et al., 2012).

Here, we have used a bioinformatics approach to search for mitochondrial proteins with an acetyltransferase catalytic domain. This analysis led to the identification of a novel splice variant of elongator protein 3 (ELP3), also known as KAT9, the known catalytic subunit of the six- subunits (Elp1–Elp6) transcription elongator complex. ELP3 has two putative enzymatic domains: the radical S-adenosylmethionine (SAM) domain contains iron-sulfur [4Fe–4S], a cluster important for the methylation of tRNAs by reductive SAM cleavage activity (Paraskevopoulou et al., 2006) (Grove et al., 2011) (Faculty et al., 2016) (Schwalm et al., 2016) and a C-terminal KAT domain with sequence similarity to the superfamily of GCN5-like acetyl transferases. Thus, ELP3 might acetylate histones (Winkler et al., 2002) and other protein targets, using acetyl-CoA as a cofactor (Y. P. Wang et al., 2014). However, ELP3 may also modify cytoplasmic (cyt)-tRNAs (Huang et al., 2005)(Lin et al., 2019). In this respect, ELP3 is required to form the 5-methoxy- carbonylmethyl-2-thiouridine (mcm5s2U34) and 5-carbamoyl-methyl-2-thiouridine (ncm5s2U34) modification of the wobble uridines U34 (position 34) in many cyt-tRNAs targets (Huang et al., 2005)(Karlsborn et al., 2014)(Lin et al., 2019). These post-transcriptional tRNA modifications are necessary for the accuracy and efficiency of protein translation (Zinshteyn & Gilbert, 2013)(Manickam et al., 2015), and the lack of them leads to variety of human diseases (Torres et al., 2014)(Hawer et al., 2019). Notably, dysregulation of ELP3’s tRNA modification activity is mainly associated with neurological disorders (Bento-Abreu et al., 2018) (Laguesse et al., 2015).

In the present study, we characterize this novel splice variant of ELP3 (mt-ELP3) and provide evidence that mt-ELP3, but not other variants, localizes to the mitochondria. Unexpectedly, mt- ELP3 is not involved in mitochondrial protein acetylation but in the post-transcriptional modification of mitochondrial (mt)-tRNAs. We show that overexpression of mt-ELP3 leads to protection of mt-tRNAs against the tRNA-specific RNase angiogenin and improves mitochondrial function by increasing mitochondrial translation, expression of oxidative phosphorylation system (OXPHOS) complexes, and mitochondrial respiration. Notably, mt-ELP3 is highly expressed in the brain, suggesting that it may play an important role in this tissue. Overall, our findings identify mt-ELP3 as a non-canonical mt-tRNA-modifying enzyme.

## Results

### Identification of ELP3 as a putative mitochondrial acetyltransferase

To identify novel mitochondrial KATs, we searched for KAT, NAT (N-acetyltransferase), and NAA (N-alpha acetyltransferase) splice products in the Amigo (http://amigo.geneontology.org/amigopaap), Panther (http://www.pantherdb.orgke), and Kegg (https://www.genome.jp/kegg/pathway.html) pathway databases and (Yang, 2004). This analysis led to the identification of 55 genes encoding a putative acetyltransferase domain in the human genome. Using the Ensemble database, we extracted 276 spliced isoforms for all candidates and we performed *in silico* mitochondrial targeting analysis using TargetP (https://services.healthtech.dtu.dk/service.php?TargetP-2.0), Predotar (http://urgi.versailles.inra.fr/predotar/predotar.html), and Wolfpsort (https://wolfpsort.hgc.jp) databases. Each spliced isoform was arbitrarily scored (the strategy is illustrated in Figure S1). The top 20 potential mitochondrial KAT isoforms include 3 splice variants of ELP3 (Table 1 and Figure S2): ELP3/Kat9 203 spliced isoform (ENSP00000439242), ELP3/Kat9 202 spliced isoform (ENSP00000445558) and ELP3/Kat9 003 spliced isoform (ENSP00000429180). Notably, the spliced isoform 202 and 203 have the same start codon (Figure S2), therefore, we considered spliced isoform 203 and 003 for subsequent studies. Subsequent studies focused on the mitochondrial localization of these isoforms and their function in mitochondria.

**Table 1.**
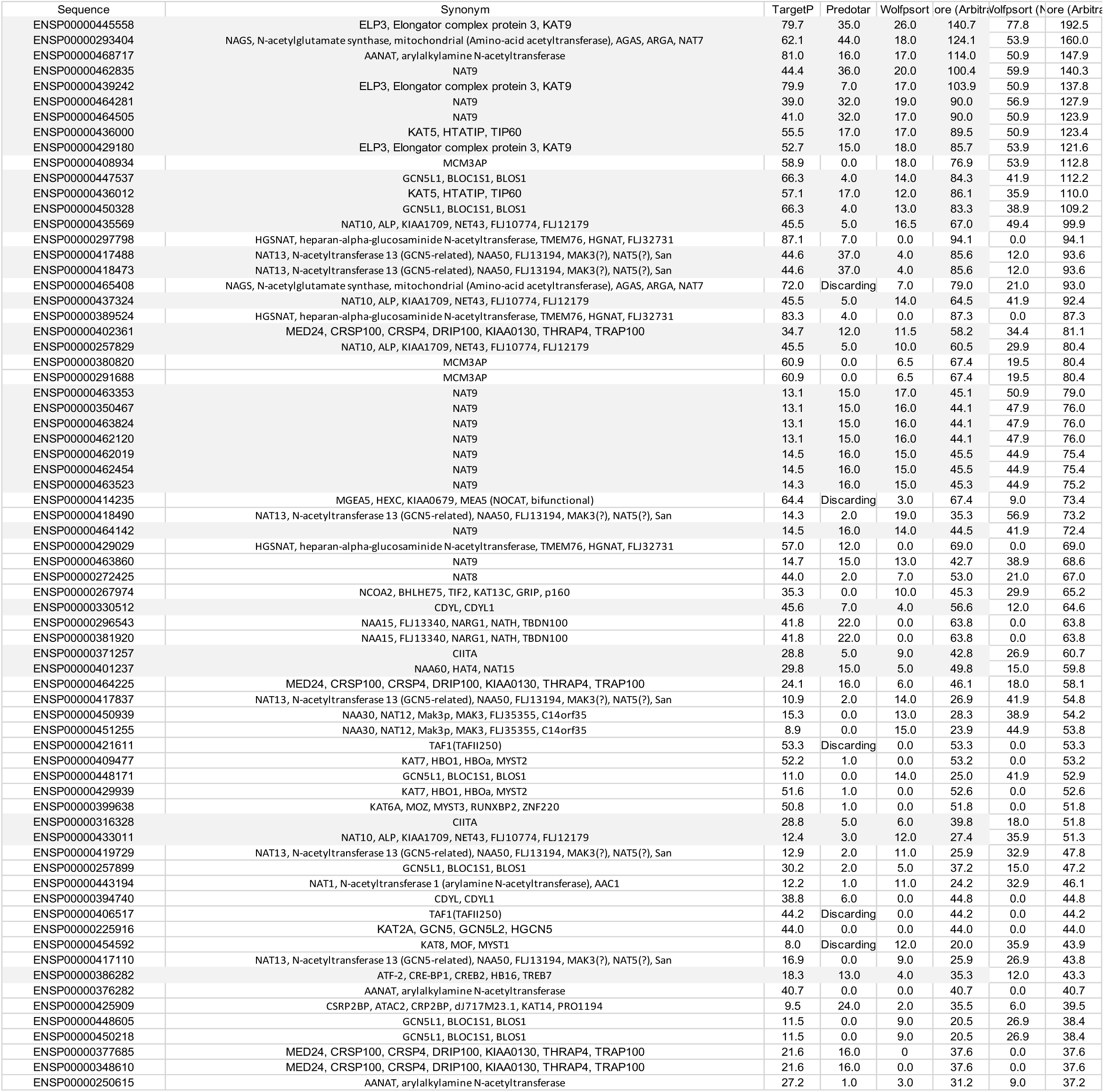
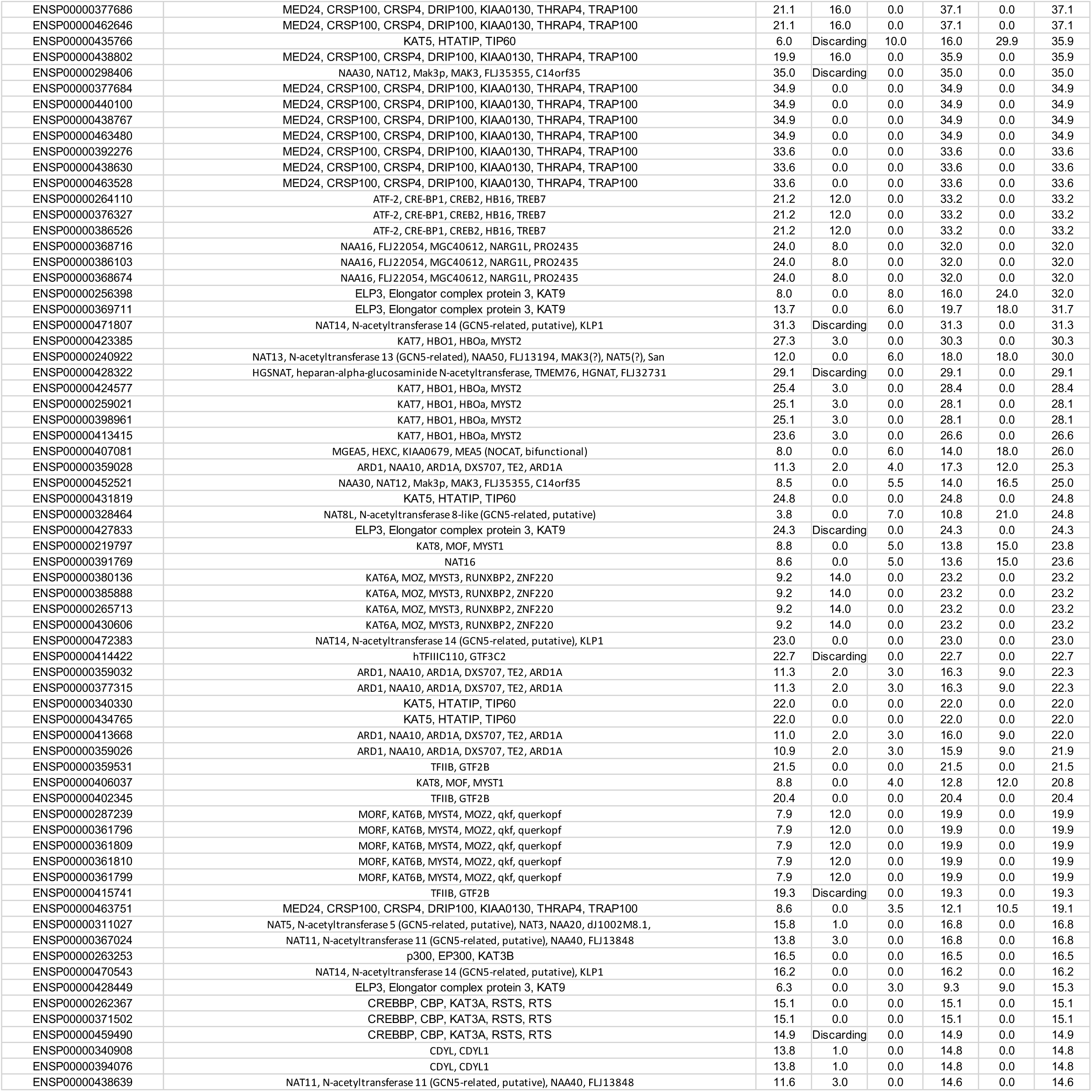
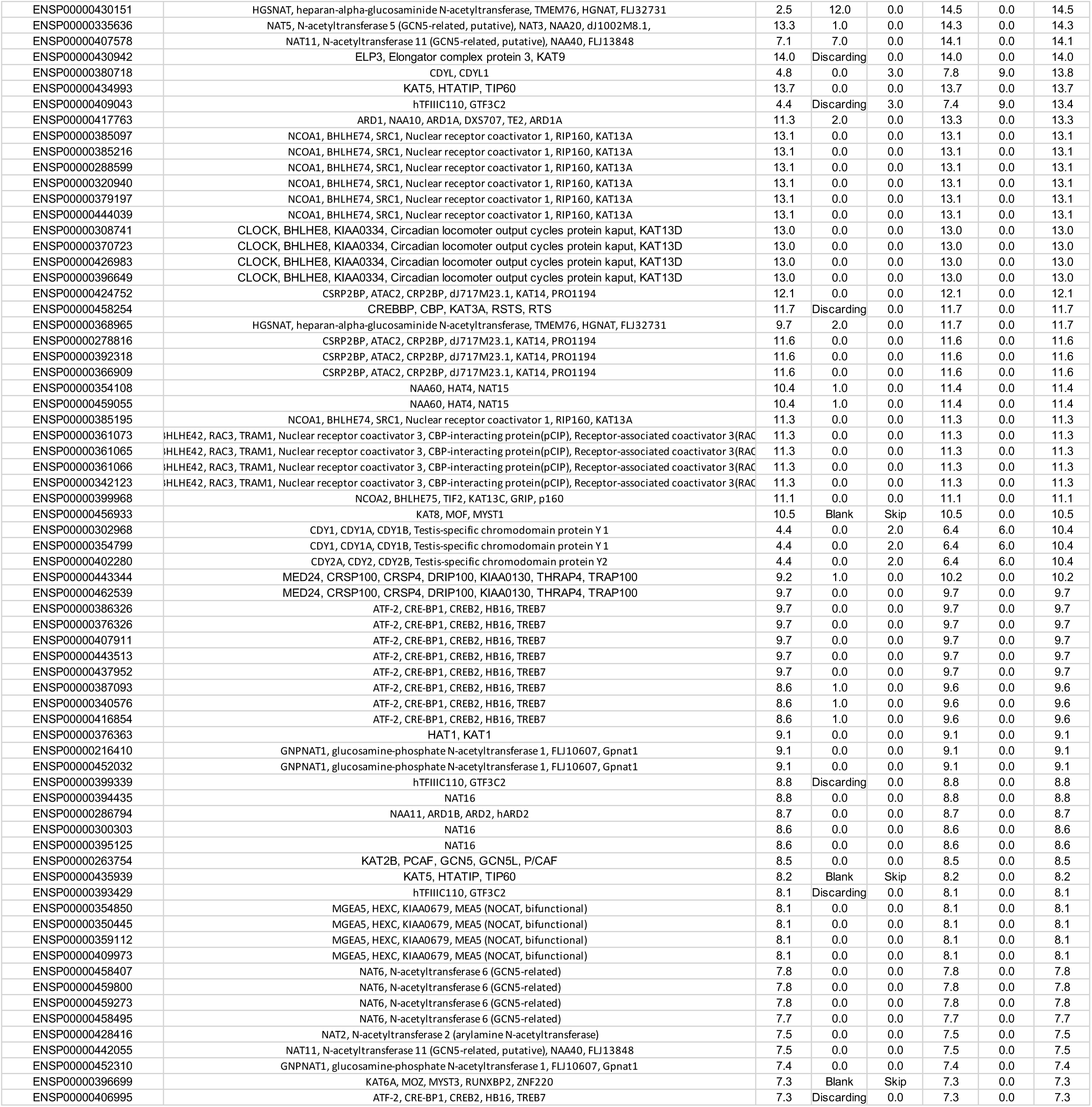
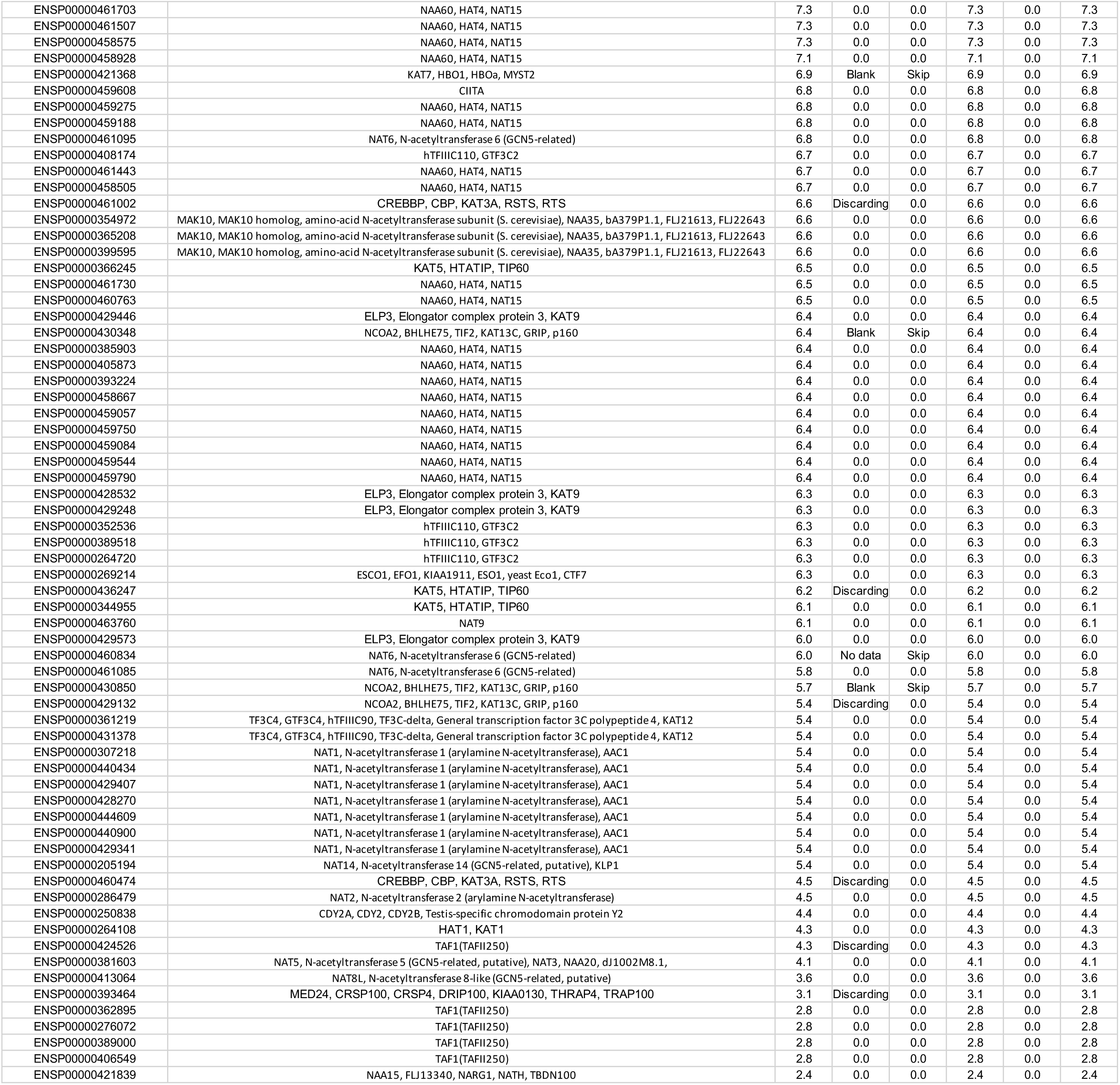
Putative mitochondrial acetyltransferases

### ELP3-203 isoform is a mitochondrial matrix protein

ELP3 is located predominantly in the cytoplasm (Pokholok et al., 2002)(Creppe et al., 2009)(Hawkes et al., 2002), but it may also localize to the mitochondria (Stilger & Sullivan, 2013)(Barton et al., 2009). To determine the subcellular localization of ELP3, canonical ELP3 (ELP3-ORF1), spliced isoform 003 (ELP3-ORF2), and spliced isoform 203 (ELP3-ORF3) (Figure S2) were C-terminally tagged with a FLAG epitope in pcDNA T0 vector and expressed transiently in HeLa cells using pcDNA T0. Immunofluorescence analysis showed that only FLAG-tagged ELP3-ORF3 co-localized with the mitochondrial marker, cox IV (Figure 1A), confirming that only this isoform (hereafter referred to as mt-ELP3) localizes to mitochondria. To investigate the sub- mitochondrial localization of mt-ELP3, we used a proteinase K sensitivity assay. A tetracycline- regulated expression system was used for inducible mt-ELP3 protein expression in HEK293T cells, and isolated mitochondria from these cells (Figure 1B) were treated with increasing concentrations of digitonin (mild detergent) in the presence of proteinase K. Mt-ELP3 was protected from cleavage by proteinase K except in high concentrations of digitonin, which dissolves the mitochondrial membranes (Figure 1C). This sensitivity was similar to that of the matrix protein HSP60 (Figure 1C). In contrast, TOM20, a mitochondrial outer membrane protein, was sensitive to proteinase K even in the absence of digitonin (Figure 1C). These findings indicate that mt-ELP3 is a mitochondrial matrix protein.

**Figure 1.**
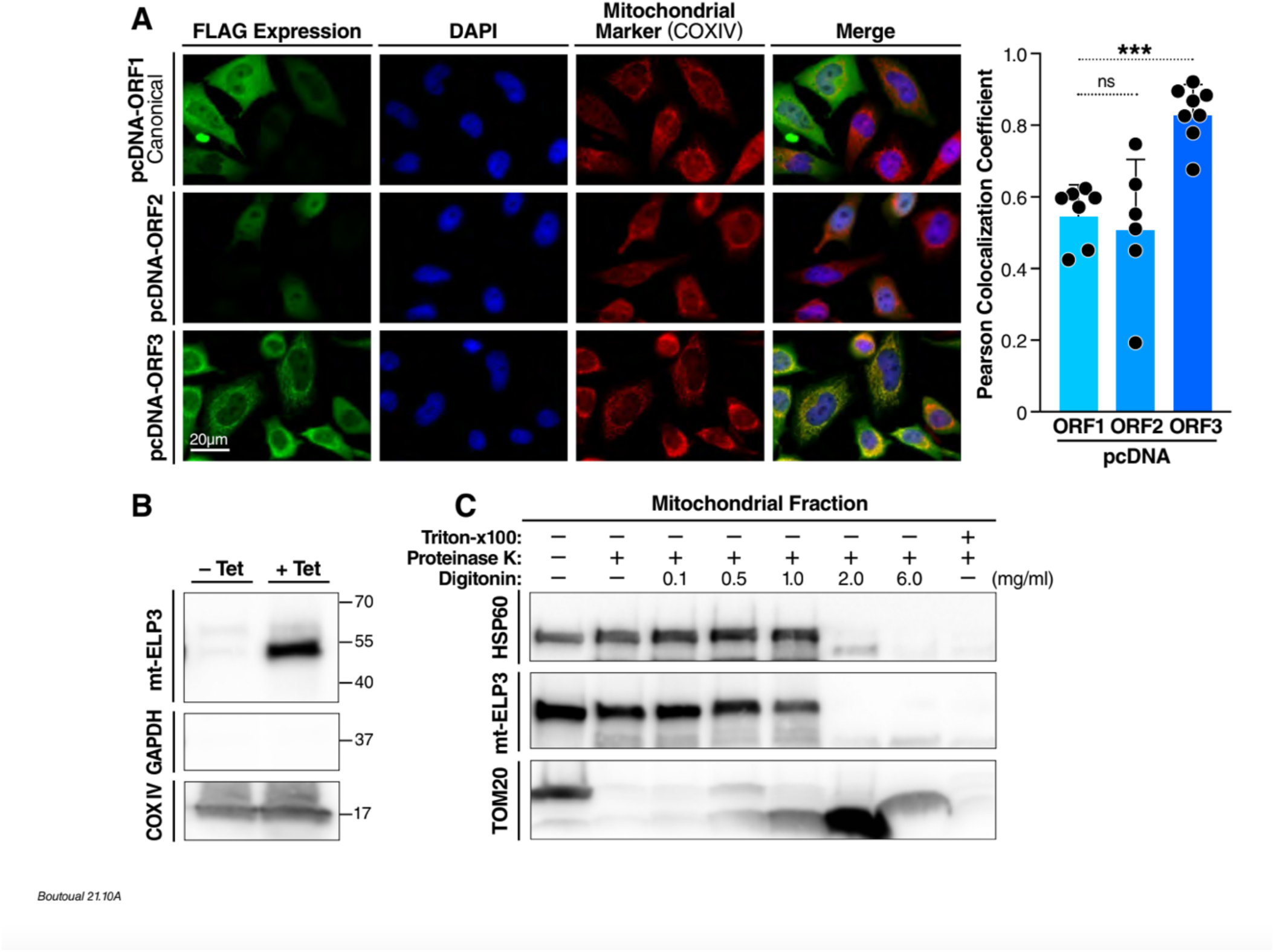
ELP3-203 isoform is a mitochondrial matrix protein. **(A)** Subcellular localization of ELP3 in mitochondria. HeLa cell lines were transiently transfected with pcDNA ELP3-ORF1, pcDNA ELP3-ORF2 and pcDNA ELP3-ORF3 plasmids. Intracellular distribution of ELP3-ORF1, ELP3-ORF2 and ELP3-ORF3 proteins was revealed with a monoclonal primary antibody against Flag and Alexa Fluor-488 (green fluorescence). Expression of COXIV (mitochondrial marker) in the same cells was assessed with a monoclonal antibody against COXIV and Alexa Fluor 594 (red fluorescence). Right panels show merged images for COXIV dye and endogenously expressed protein ELP3-ORF1, ELP3-ORF2 and ELP3-ORF3 in HeLa cells. Yellow color shows colocalization of COXIV with ELP3-ORF1, ELP3-ORF2 and ELP3-ORF3. Nuclei (blue) were visualized with DAPI. These images are representative of three separate experiments. The accompanying histogram shows quantitative analysis of ELP3-ORF1, ELP3-ORF2 and ELP3-ORF3 co-localized with COXIV marker in HeLa cells by Pearson’s correlation coefficient. All data are mean ± SD of at least three experiments. ***p < 0.001. n.s.: not significant. **(B)** Western blot analysis of mt-ELP3 protein in the mitochondrial fraction from T-REx-293 cells treated (+Tet) or not (-Tet) with tetracycline. GAPDH and COXIV were used as cytosolic and mitochondrial protein markers, respectively. **(C)** Proteinase K sensitivity of mt-ELP3. Purified mitochondria from cells overexpressing mt- ELP3 were left untreated (-), treated (+) with proteinase K, or treated with proteinase K in the presence of increasing concentrations of digitonin, or proteinase K with 0.3% Triton X-100. HSP60 and TOM20 were used as mitochondrial mitoplast and outer membrane markers, respectively.

### Overexpression of mt-ELP3 does not affect mitochondrial protein acetylation

ELP3 contains a histone/lysine acetyltransferase domain in its carboxy-terminus with significant homology to the catalytic domain of the GNAT (Gcn5-related N-terminal acetyltransferase) family of acetyltransferases (Winkler et al., 2002). ELP3 acetylates histones (Winkler et al., 2002) and other non-histones proteins (Y. P. Wang et al., 2014)(Creppe et al., 2009). Protein lysine acetylation is an important regulatory mechanism of metabolic enzymes in the mitochondria (Bjoern Schwer et al., 2006)(Parodi-Rullán et al., 2018)(Baeza et al., 2016).

Since mt-ELP3 has an acetyltransferase domain, we speculated that it may affect the acetylation status of mitochondrial proteins. To test this hypothesis, we evaluated global mitochondrial acetylation levels in control cells and cells overexpressing mt-ELP3 using two different methods: western blot and mass spectrometry. Western blot analysis with a commercial pan-acetyllysine antibody revealed no global change of mitochondrial protein acetylation levels in cells overexpressing mt-ELP3 compared to control cells (Figure 2). Mass spectrometry analysis also showed no global change of mitochondrial acetylation level in cells overexpressing mt-ELP3, compared to control cells (Table 2).

**Figure 2.**
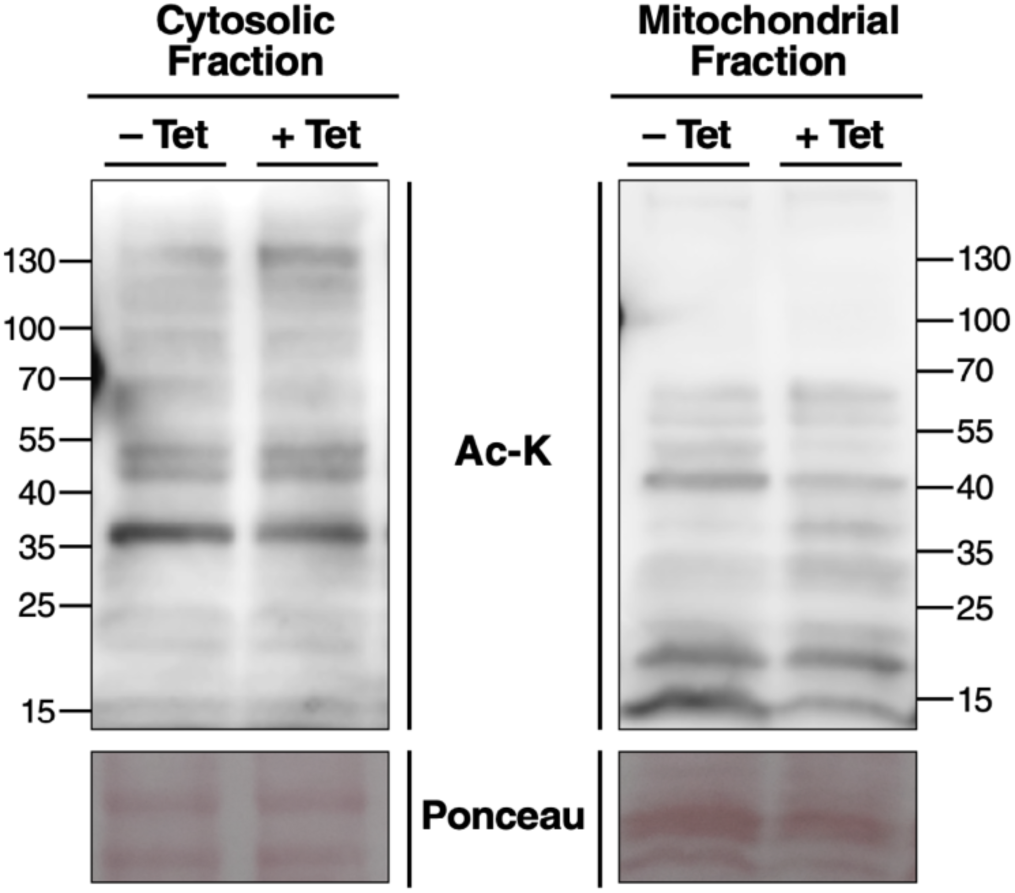
Overexpression of mt-ELP3 does not affect mitochondrial protein acetylation. Western blot analysis of lysine acetylation (Ac-K) status in mitochondrial and cytoplasmic fraction isolated from T-REx-293 cells treated (+Tet) or not (-Tet) with tetracycline. Ponceau was used as a loading control.

**Table 2.**
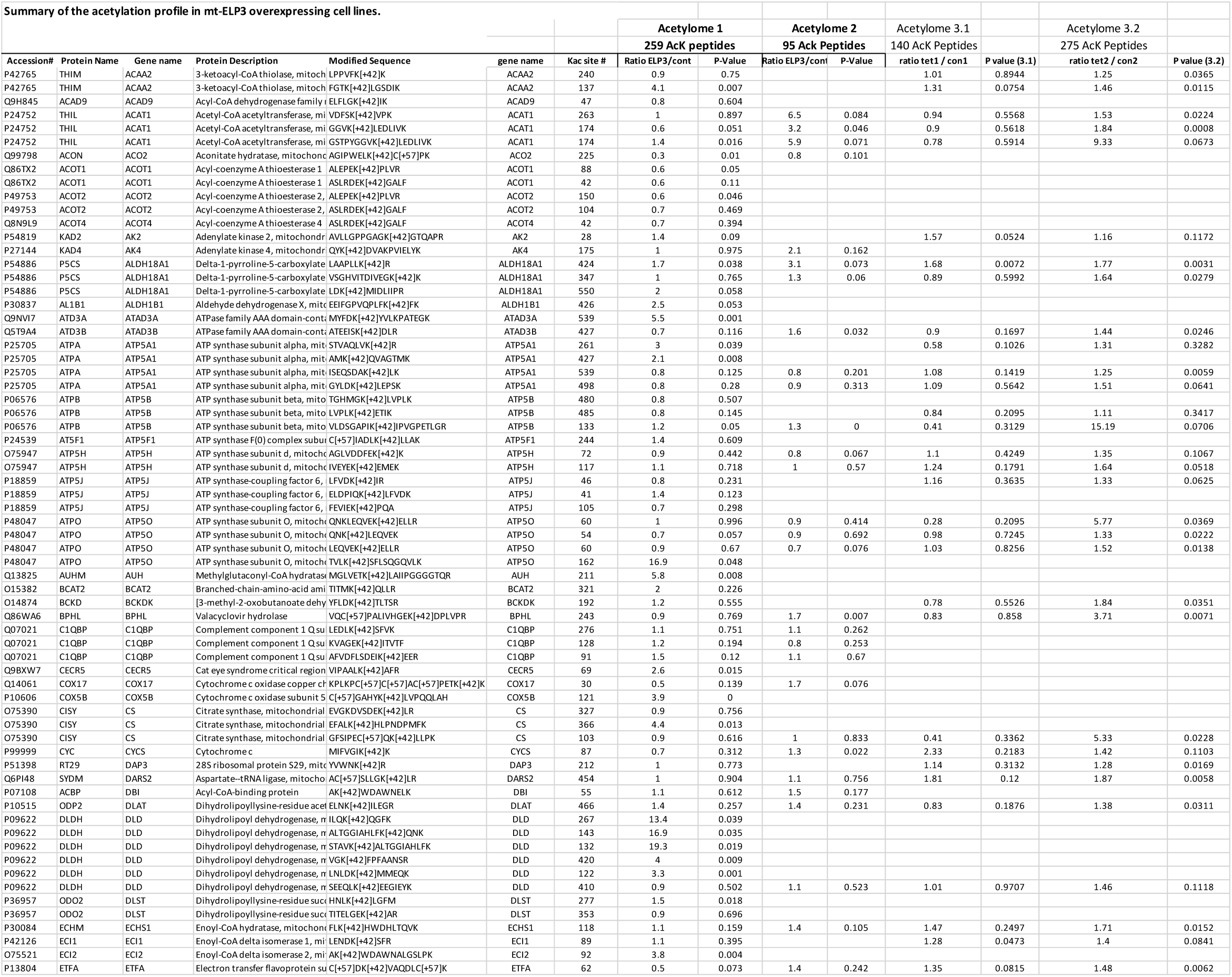
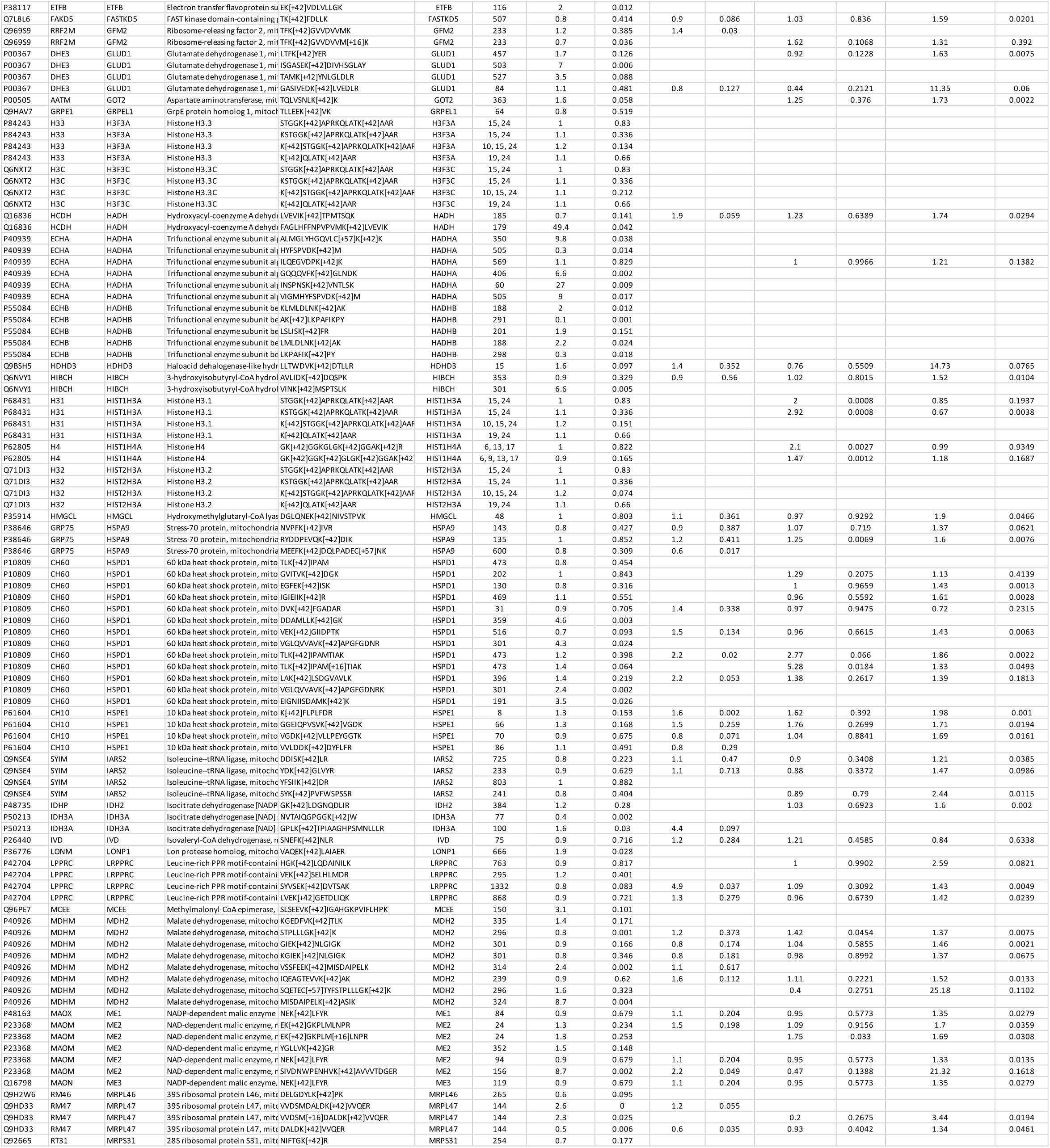
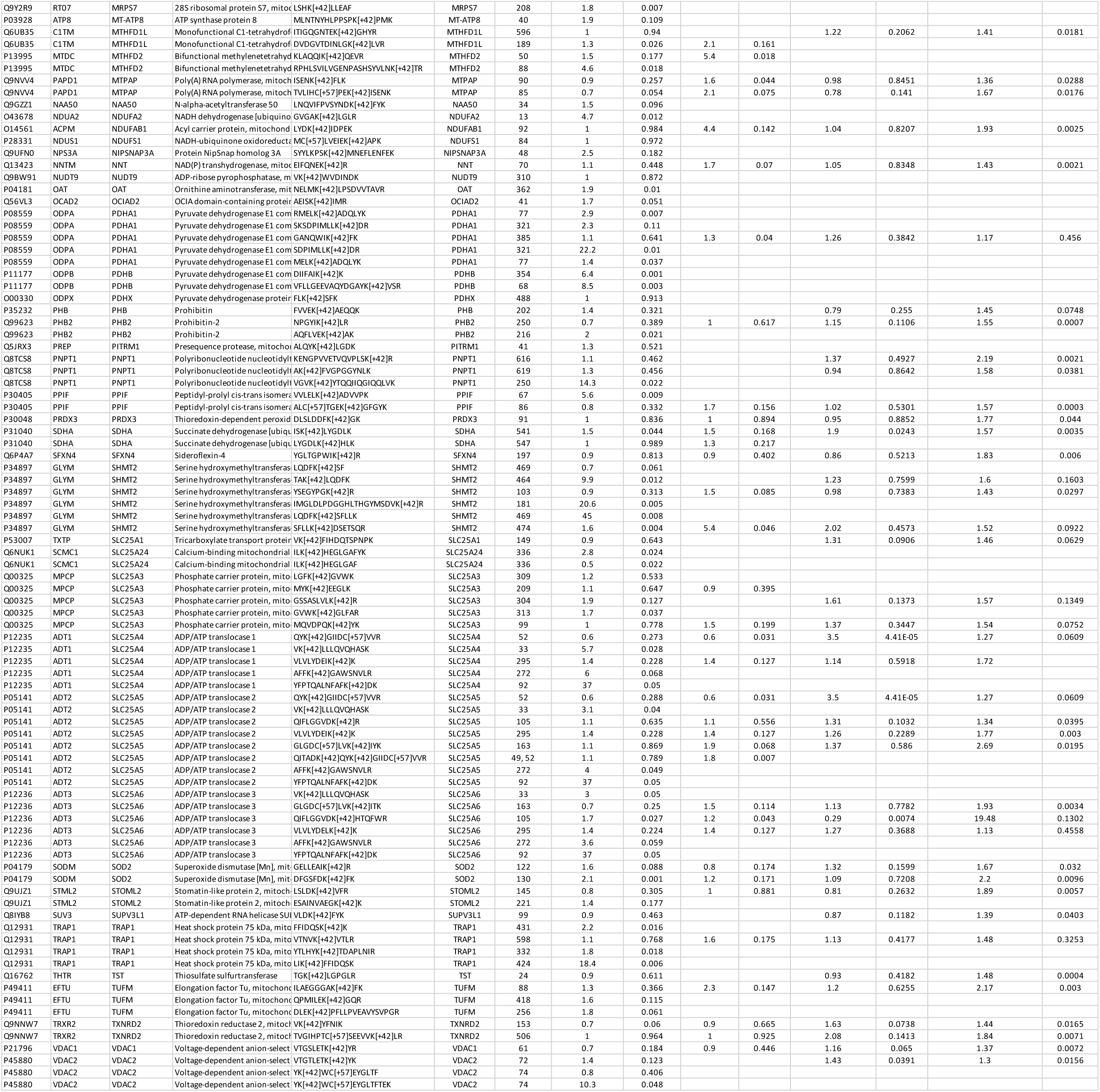
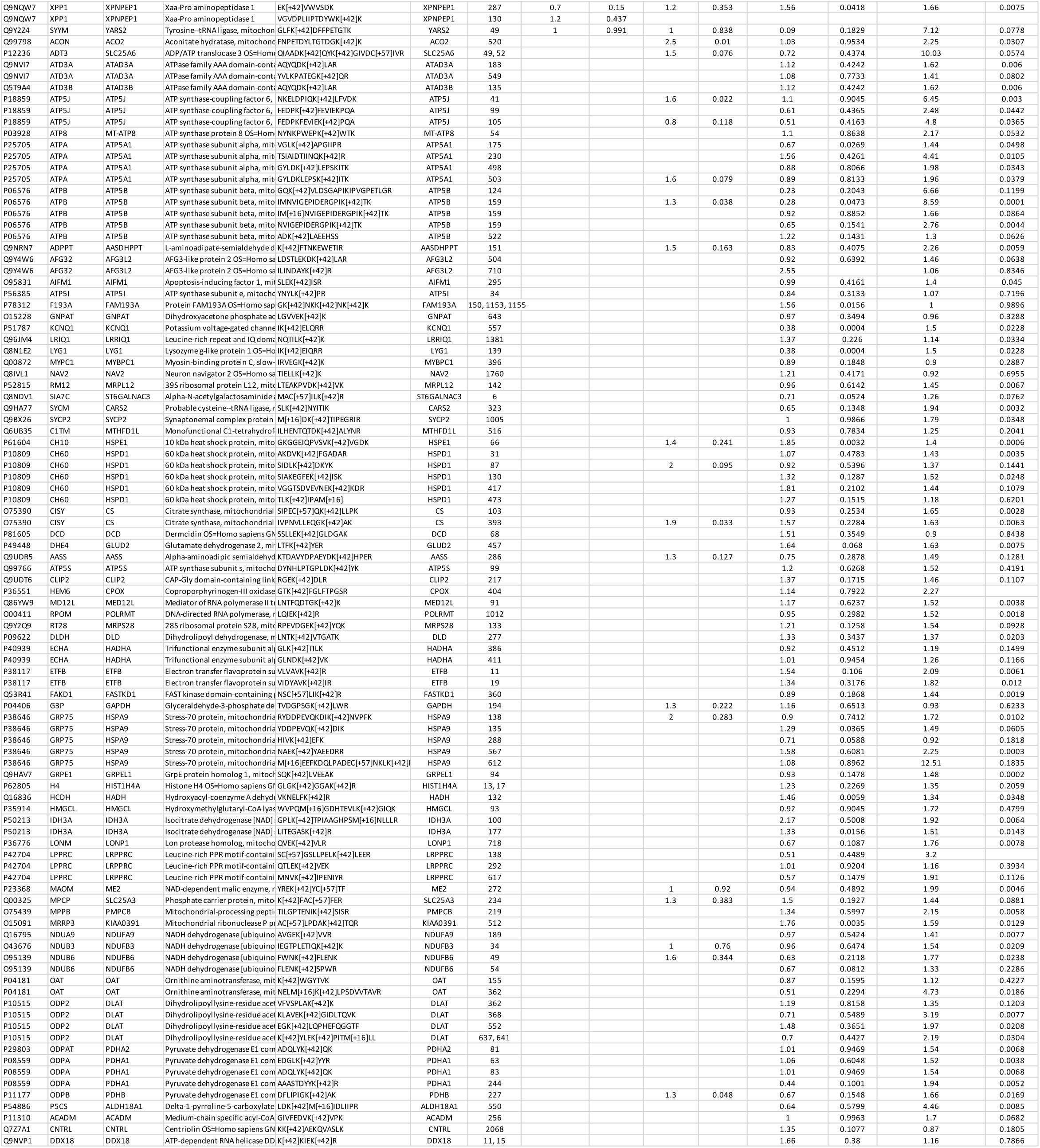
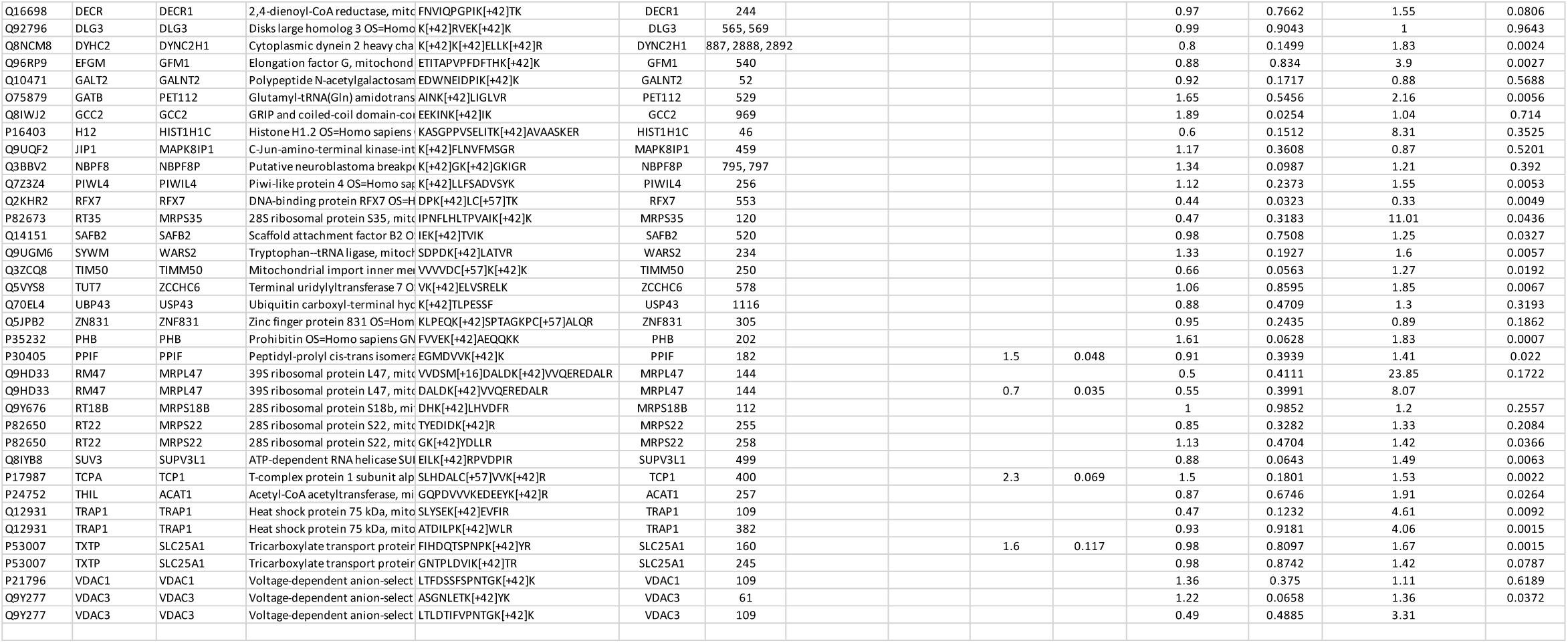
Summary of the acetylation profile in mt-ELP3 overexpressing cell lines.

### Overexpression of mt-ELP3 protects mt-tRNAs from angiogenin-mediated cleavage without altering 2-thiolation levels

Next, we tested whether mt-ELP3 is involved in mt-tRNA modifications. ELP3 not only catalyzes the acetylation of lysine residues in peptides and proteins, it also acts as a non-canonical tRNA modifying enzyme (Dauden et al., 2019)(Lin et al., 2019). In eukaryotes, canonical ELP3 is required for synthesis of 5-methoxycarbonylmethyl (mcm5) and 5-carbamoylmethyl (ncm5) side chains on uridines at the wobble position (position 34 in the tRNA) of a subset of cyt-tRNAs (Huang et al., 2005). The mcm^5^U_34_-modified wobble base is further thiolated to 5-methoxycarbonylmethyl-2-thiouridine (mcm^5^_S_^2^U) in tRNA^Lys^_UUU_, tRNA^Glu^_UUC_ and tRNA^Gln^_UUG_ (Leidel et al., 2009), and the presence of a mcm5 or ncm5 side chain is a prerequisite for efficient thiolation in these cyt-tRNAs (C. Chen et al., 2011). Corresponding mt-tRNAs are also modified at the wobble uridine position, which is 5-taurinomethyl-2-thiouridine (τm^5 2^U) (Suzuki & Suzuki, 2014).

To evaluate a possible role of mt-ELP3 in modifying mt-tRNAs, we analyzed the sensitivity to digestion with the tRNA-specific RNase angiogenin of mt-tRNA^Glu^, mt-tRNA^Lys^ and mt-tRNA^Leu^ (with the τm5U modification) and mt-tRNA^Val^ (without the τm5U modification) from control cells and from cells overexpressing mt-ELP3. tRNAs lacking modification at the wobble position are more sensitive toward angiogenin-mediated digestion than the corresponding modified tRNAs (Lefkimmiatis et al., 2019) (Navarro-González et al., 2017)(R. Boutoual et al., 2018). After *in vitro* angiogenin digestion of total RNA purified from control cells and cells overexpressing mt-ELP3, the digested products were analyzed using northern blot analysis with a specific digoxigenin (DIG)-labeled oligodeoxynucleotide probe for the mentioned mt-tRNAs. Mt-tRNA^Glu^, mt- tRNA^Lys^ and mt-tRNA^Leu(UUR)^ from control cells were more sensitive to angiogenin-mediated cleavage than those from cells overexpressing mt-ELP3 (Figure 3A). In contrast, we found no differences in the digestion patterns of mt-tRNA^Val^ (which carries no τm^5^U modification) (Figure 3A). These results are consistent with the model that mt-ELP3 increases the synthesis or stability of τm^5^U modification in these mt-tRNAs.

**Figure 3.**
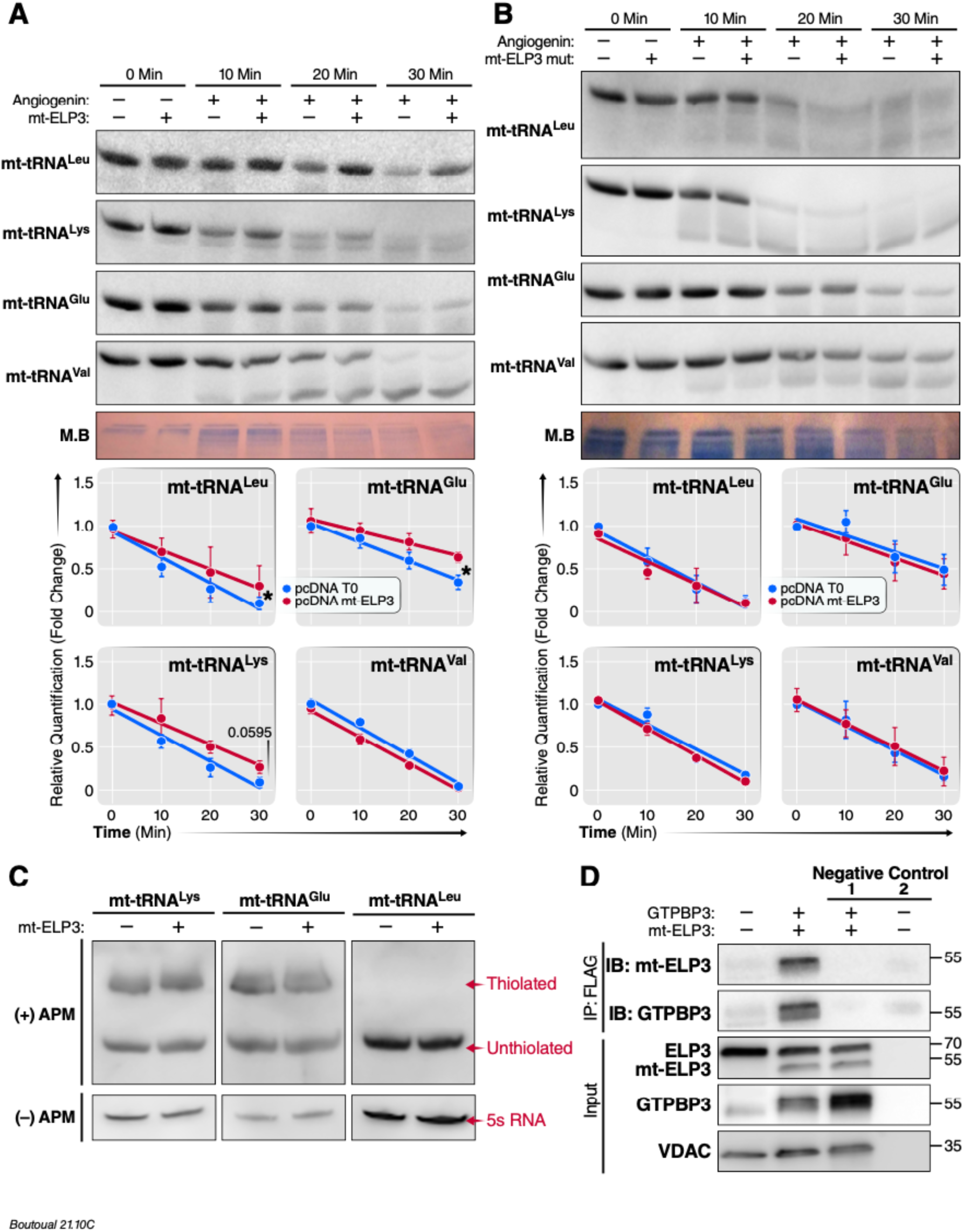
Overexpression of mt-ELP3 protects mt-tRNAs substrate from angiogenin-mediated cleavage without altering 2-thiolation levels **(A and B)** Northern analysis of mt-tRNA^Lys,^ mt-tRNA^Leu^, mt-tRNA^Glu^ and mt-tRNA^Val^ after in vitro angiogenin digestion of small RNAs purified from control cells and cells overexpressing mt- ELP3 (**A**), and from control cells and cells overexpressing mt-ELP3 mut (**B**) for the indicated times. Total RNA stained with methylene blue (MB) was used as loading control. The accompanying histogram shows the kinetic analysis of angiogenin digestions. The amounts of intact mt-tRNA after 0, 10, 20 and 30 min of incubation with angiogenin are represented as fold-changes relative to the undigested control (0 min). All data are mean ± SD of at least three experiments. *p < 0.05. **(C)** APM-northern analysis of 2-thiolation of mt-tRNA^Lys^, mt-tRNA^Leu^ and mt-tRNA^Glu^ from control cells and cells overexpressing mt-ELP3. The same amounts of total RNA (5 µg) were run in a denaturing polyacrylamide-urea gel in the presence (+) or absence (−) of APM. The thiolated tRNAs were detected as retarded bands in the presence of APM. The APM (−) membrane was probed with 5S rRNA as a loading control. **(D)** mt-ELP3 interacts with GTPBP3. HEK 293T cells were co-transfected with pcDNA mt-ELP3- Flag and pCMV6 GTPBP3 plasmids as indicated, and the lysates were immunoprecipitated with anti-Flag antibody. Precipitated proteins were immunoblotted with anti-ELP3, GTPBP3 and VDAC antibodies. Beads mixed with lysate without anti-Flag antibody and beads mixed with anti-Flag antibody without lysate were used as a negative control 1 and 2, respectively.

To test whether the KAT domain of mt-ELP3 is necessary for mt-tRNA modification, we generated a mutated mt-ELP3 construct (mt-ELP3 mut) by introducing a (GFG→FAF) mutation in its KAT domain (Selvadurai et al., 2014) (Padgett et al., 2018)(Figure S3-4). We found no differences in the angiogenin-mediated cleavage of RNA from control cells and cells overexpressing mt-ELP3 mut (Figure 3B), indicating that the KAT domain of mt-ELP3 is essential for its mt-tRNAs function.

The τm^5^U_34_ modification is further thiolated to τm^5^_S_ U in mt-tRNA^Lys^, mt-tRNA^Glu^ and mt-tRNA^Gln^ (Boczonadi et al., 2013)(Wu et al., 2016). We asked whether mt-ELP3 also affects 2- thiouridylation in these mt-tRNAs. Thiolation of U34 can be detected by northern blot in the presence of [(N-acryloylamino)phenyl]mercuric chloride (APM). In this analysis, the APM polymerized in the gel specifically interact with tRNAs containing thiol group, thereby retarding their migration (Lemau de Talancé et al., 2011)(X. Wang et al., 2010). 2-Thiouridylation levels of mt-tRNA^Glu^ and mt-tRNA^Lys^ in cells overexpressing mt-ELP3 were comparable with those in control cells (Figure 3C). In contrast, no retardation was observed with mt-tRNA^Leu^, which is natively non-thiolated. These observations indicate that mt-ELP3 does not affect thiolation at position 2 of mt-tRNAs.

Overexpression (Figure S5A) or down-regulation (Figure S5B) of canonical ELP3 leads to an increase and decrease of 2-thiol level, respectively, in cyt-tRNAs (Figure S5 C-D); however, overexpression of mt-ELP3 did not increase thiol level in mt-tRNAs (Figure 3C), supporting the idea that both τm^5^ and 2-thiol modifications in mt-tRNAs appear to proceed independently, similarly to published reports (Umeda et al., 2005) (Martínez-Zamora et al., 2015). The nuclear-encoded tRNA-modifying enzymes, MTO1 and GTPBP3, are thought to be responsible for the synthesis of τm^5^U34 modification in mt-tRNAs encoding for Lys, Leu, Glu, Gln and Trp (Asano et al., 2018). Based on our observations that mt-ELP3 seems to be involved in the synthesis of τm^5^U, we predicted that mt-ELP3 interact with GTPBP3 as well as MTO1. Mass spectrometry analysis of proteins that co-immunoprecipitate with mt-ELP3-FLAG led to the identification of GTPB3 (data not shown). To confirm this interaction, we co-expressed GTPBP3 and Flag-tagged mt-ELP3 in HEK293T cells and we found that by pulling down with a Flag-tag antibody, GTPBP3 could be detected (Figure 3D). These results confirm the protein–protein interaction between mt- ELP3 and GTPBP3.

### Overexpression of mt-ELP3 results in an increase in mitochondrial translation

A taurine modification at U34 (τm^5^_S_ U34) is crucial for regulating mitochondrial translation efficiency and accuracy of codon-anticodon interaction (Kirino et al., 2005), and the GTPBP3 defect causes a deficiency in mitochondrial translation (Martínez-Zamora et al., 2015) (D. Chen et al., 2019). Thus, we reasoned that overexpressing mt-ELP3 might increase mitochondrial translation. To examine mitochondrial translation activity, we used a non-radioactive pulse- labelling click chemistry experiment (Zorkau et al., 2020). Specifically, we treated control cells and cells overexpressing mt-ELP3 with the alkyne-methionine derivative, L- homopropargylglycine (HPG), with or without cycloheximide (specific inhibitor of cytosolic translation) and/or chloramphenicol (a specific inhibitor of mitochondrial translation). The methionine analogue HPG was incorporated into nascent protein in the place of methionine and subsequently visualized using chemoselective fluorescence-tagging by means of click chemistry (Dieterich et al., 2010)(Zorkau et al., 2020).

As predicted, mitochondrial translation was increased in cells overexpressing mt-ELP3 compared to control cells (Figure 4A, middle right and 4B (HPG+CHX)). Notably, the increase was eliminated by chloramphenicol treatment, indicating that cycloheximide-resistant signal represents exclusively newly synthesized mitochondrial proteins (Figure 4A, right and 4B (HPG+CHX+CHL)). However, no signal was detected without HPG (Figure 4A, left). Furthermore, the total amount of newly synthesized proteins (cytosolic and mitochondrial) was greater in cells overexpressing mt-ELP3 than in control cells as evidenced by HPG signal without cycloheximide and chloramphenicol (Figure 4A, middle left and 4B (HPG)). As a support to the HPG signal measurements, we also quantified the HPG signal in control cells and cells overexpressing mt-ELP3 with cycloheximide (mitochondrial translation) by flow cytometry. We found significantly more HPG signal in cells overexpressing mt-ELP3 than control cells, consistent with the data from fluorescence microscopy (Figure 4C and D). These observations indicate that overexpression of mt-ELP3 increases the mitochondrial translation rate.

**Figure 4.**
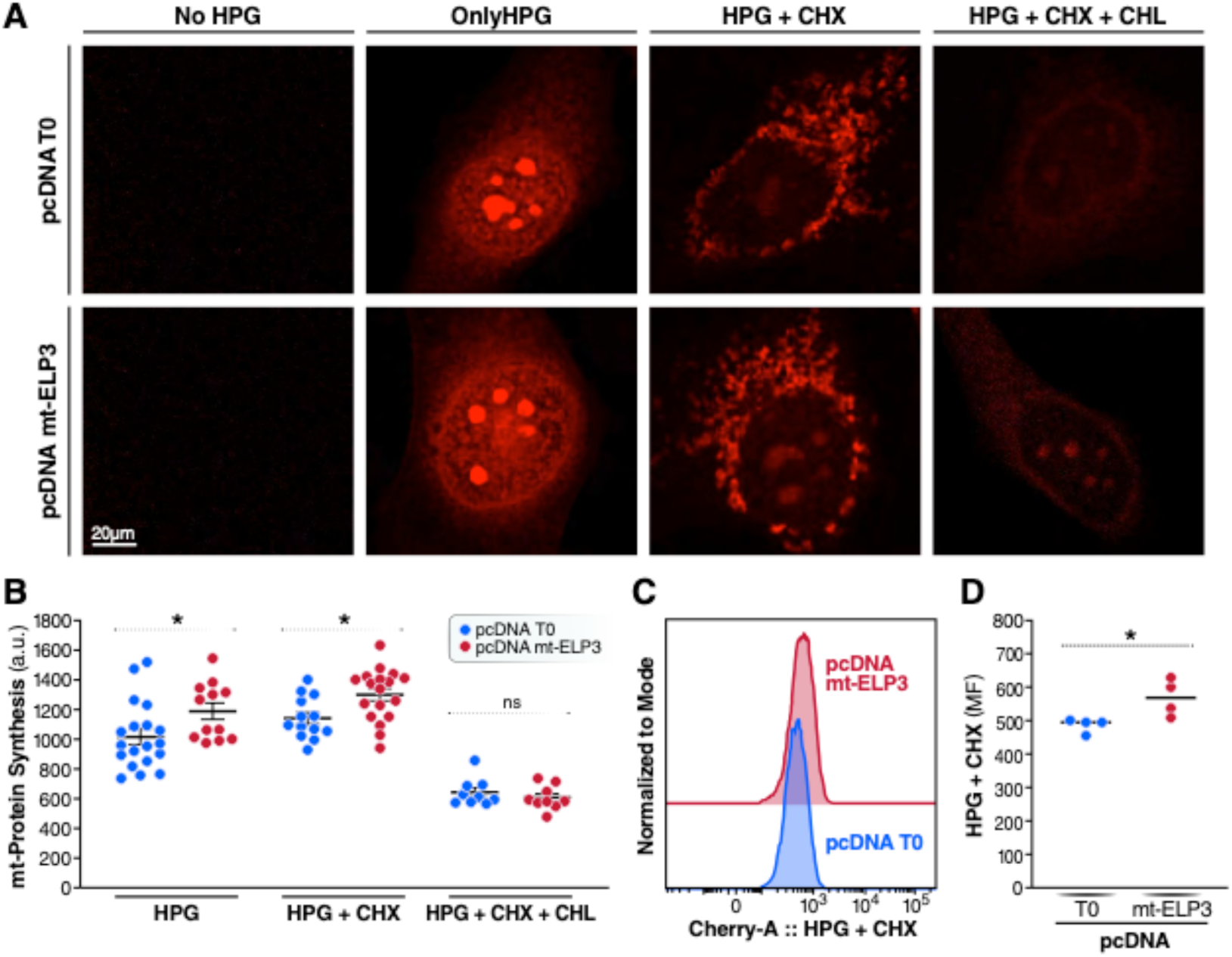
Overexpression of mt-ELP3 results in an increase in mitochondrial translation. **(A, B, C, and D)** Evaluation of mitochondrial translation. De novo mitochondrial protein synthesis was visualized in HEK293T cells transiently transfected with empty plasmid (pcDNA T0) or mt- ELP3 plasmid (pcDNA mt-ELP3) by incubation with or without HPG (1h) and with or without cycloheximide (CHX) and/or chloramphenicol (CHL). After 1h, the cells were fixed, underwent the click reaction and analyzed for the flurorescence-tagged HPG incorporation by flurorescence microscopy (**A**) or flow cytometry (**C**). The HPG signal was calculated and presented graphically (**B** and **D**). All data are mean ± SD of at least three experiments. *p < 0.05. n.s.: not significant.

### Overexpression of mt-ELP3 increases the expression of mitochondrial OXPHOS complexes and mitochondrial respiration

Mitochondrial translation is responsible for the synthesis of 13 mitochondrially encoded subunits of OXPHOS. To assess a potential role on OXPHOS, we measured the steady-state levels of OXPHOS complexes by Blue-Native gel electrophoresis (Figure 5A). Interestingly, levels of complex I (C I), complex III (C III), complex IV (C IV) and complex V (C V) were greater in cells overexpressing mt-ELP3 than in control cells (Figure 5B). Importantly, overexpression of mt- ELP3 mut did not affect the expression of these complexes (Figure 5C-D). Notably, the increased expression of OXPHOS complexes in cells overexpressing mt-ELP3 was accompanied by an increase in the steady-state levels of mitochondrial-encoded OXPHOS subunits (Figure 5E). No change was observed in the levels of mitochondrial-encoded OXPHOS subunits when overexpressing mt-ELP3 mut (Figure 5E).

**Figure 5.**
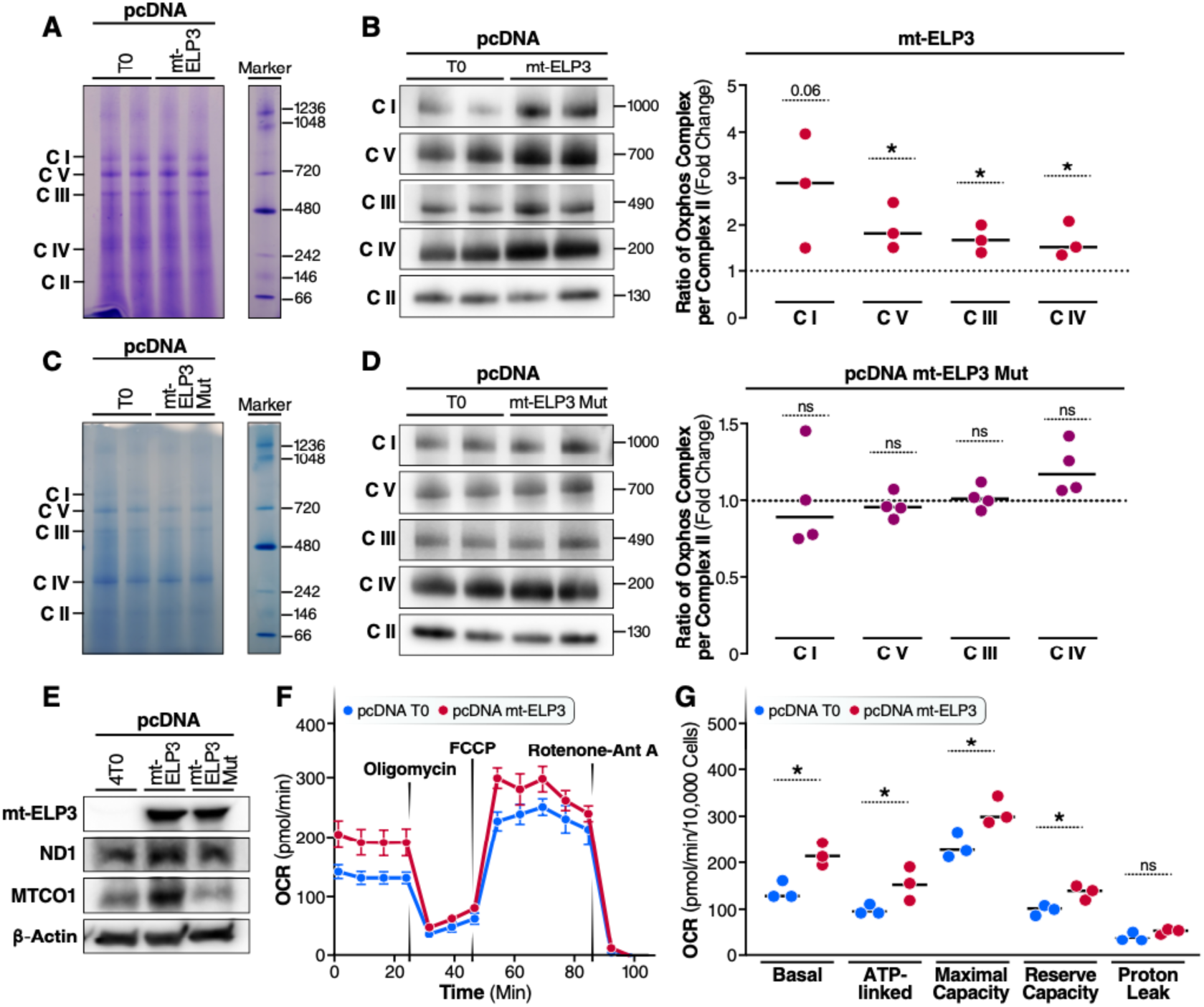
Overexpression of mt-ELP3 results in an increase in the expression of mitochondrial OXPHOS complexes and mitochondrial respiration (A and C) Representative BN-PAGE gel stained with Coomassie Brilliant Blue of respiratory complexes purified from control cells (pcDNA T0) and cells overexpressing mt-ELP3 (pcDNA mt-ELP3) (A), and from control cells (pcDNA T0) and cells overexpressing mt-ELP3 mutant (pcDNA mt-ELP3 mut) (C). A 20-µg aliquot of each sample was loaded per well. A 5-µL aliquot of NativeMark unstained protein standard (Marker) was loaded. (B and D) Representative Blue Native-PAGE of OXPHOS complexes in HEK293T cells transfected with pcDNA4 T0 or pcDNA mt-ELP3 (B) and with pcDNA4 T0 or pcDNA mt-ELP3 mut (D). The scatter plots show the densitometric measurements of OXPHOS complexes normalized to complex-II (loading control) and represented as fold change relative to control cells. **(E)** Representative immunoblots of mitochondrially encoded OXPHOS subunits ND1 (Complex I), MTCO1 (Complex IV) in HEK293T cells transfected with pcDNA4 T0, pcDNA mt-ELP3 or pcDNA- mt-ELP3 mut. The membranes were also probed with an antibody against ELP3 and b-actin, B-actin was used as a loading control. **(F and G)** Oxygen consumption rate (OCR). Equal numbers of HEK293T cells transiently transfected with pcDNA T0 or pcDNA mt-ELP3 were subjected to oxygen consumption measurements in a Seahorse XF96 extracellular flux analyzer, with sequential additions of the metabolic inhibitors/activators: oligomycin, FCCP, antimycin A (Ant A) and rotenone (**F**) The scatter plot shows basal OCR (determined as the difference between OCR before oligomycin and OCR after rotenone/antimycin A/rotenone), ATP-linked OCR (difference between OCR before and after oligomycin), maximal OCR, reserve capacity (difference between the FCCP-stimulated rate and basal OCR) and proton leak (difference between basal OCR and ATP-linked OCR) in pcDNA T0 and pcDNA mt-ELP3 cells (**G**). All data are mean ± SD of at least three different experiments. *p < 0.05, **p < 0.01. n.s.: not significant.

Next, we tested the effect of mt-ELP3 overexpression on mitochondrial respiration using a Seahorse XF96 extracellular flux analyzer with a series of metabolic inhibitors and uncoupling agents (Figure 5F). We observed that the basal oxygen consumption rate (OCR), mitochondrial ATP-linked and reserve respiratory capacity (ATP that can be utilized during increases in energy demand) were significantly greater in mt-ELP3-overexpressing cells than control cells and no difference was found in ATP leak levels between these cells (Figure 5G). Overall, these findings indicate that overexpression of mt-ELP3 increase mitochondrial respiration.

## Discussion

In the present study, we attempted to identify mitochondrial acetyltransferase candidates by a bioinformatic approach. This led to the identification of a novel splice variant and isoform of ELP3 localized to the mitochondrial matrix. Surprisingly, this mitochondrial ELP3 (mt-ELP3) is not involved in the acetylation of mitochondrial proteins, but rather in the post-transcriptional modification of mt-tRNAs, whereby it regulates mitochondrial RNA translation and respiration. Our results clearly demonstrated the localization of a short form of ELP3 in the mitochondrial mitoplast. While previous reports have indicated that ELP3 localizes to mitochondria (Barton et al., 2009) (Padgett et al., 2018), an ELP3 homologue in *Toxoplasma gondii* is a tail-anchored protein located in the parasite’s outer mitochondrial membrane due to its unique C-terminal transmembrane domain (TMD)(Padgett et al., 2018). However, mammalian ELP3 lacks a TMD domain indicating that another mechanism, most likely the presence of NH2-terminal mitochondrial targeting sequences, is involved in its mitochondrial import.

The localization of ELP3 to mitochondria suggested a novel and unknown function in this organelle. Unexpectedly, overexpression of mt-ELP3 did not increase mitochondrial lysine acetylation levels. Recent studies have showed that ELP3 acts as a non-canonical acetyltransferase that catalyzes reactions on tRNAs instead of proteins (Lin et al., 2019) (Abbassi et al., 2020)(Dauden et al., 2019). Therefore, ELP3 is now considered a tRNA-modifying enzyme that catalyzes the initial step in the formation of mcm^5^U, mcm^5^s^2^U, and ncm^5^U on uridines in the wobble base position of a subset of cytosolic tRNAs (Lin et al., 2019).

We reasoned that mt-ELP3 might be involved in the post-transcriptional modification of mt-tRNAs. Like cytoplasmic tRNAs, a subset of mitochondrial tRNAs is also modified at the U34 position. A τm^5^_S_^2^ U modification is the mitochondrial counterpart of mcm^5^ s^2^ U (Suzuki & Suzuki, 2014). In agreement with the hypothesis that mt-ELP3 might modify mitochondrial tRNAs, we found that mt-ELP3 overexpression decreases the sensitivity of mt-tRNA^Leu^, mt-tRNA^Lys^ and mt- tRNA^Glu^ to angiogenin digestion, indicating a regulatory role for mt-ELP3 in modifying τm^5^_S_ U34. Noteworthy, a defect in MTO1 and/or GTPBP3, two proteins that are predicted to jointly catalyze the addition of the τm^5^_S_ U34 modification, increases the sensitivity of their mt-tRNAs substrates (mt-tRNAs decoding for Lys, Glu, Gln, Leu^(UUR)^, and Trp) to angiogenin digestion (D. Chen et al., 2019)(Navarro-González et al., 2017) (R. Boutoual et al., 2018). Furthermore, our results showed that mt-ELP3 physically interacts with GTPBP3. Therefore, we conclude that mt-ELP3 has a role in the biogenesis of the τm^5^_S_ U modification of these mt-tRNAs.

Our data also indicate that mt-ELP3 does not control thiol modification at position 2 since overexpression of mt-ELP3 did not affect the thiolation status of these mt-tRNAs. These data further indicate that the thiolation at position 2 (^2^U34) is independent of the modification at position 5 (τm^5^U34) (Navarro-González et al., 2017) (Martínez-Zamora et al., 2015). As noted above, MTO1 and GTPBP3 cooperatively catalyze the τm^5^U34 modification since a defect in these proteins directly results in deficient τm^5^U34 modification of their mt-tRNA substrates. However, Asano et al. (2018) showed that the formation of τm^5^U34 had an efficiency of only 3.3% when attempting to reconstitute τm^5^U34 in vitro with the GTPBP3-MTO1 complex, taurine, 5,10-CH2- THF, GTP and co-factors (Asano et al., 2018), suggesting that an uncharacterized protein is necessary for efficient τm^5^U34 formation. We suggest that mt-ELP3 plays a role in the biogenesis of τm^5^U34, and further studies evaluating τm^5^U34 formation in the presence of mt-ELP3 are warranted.

Alternatively, mt-ELP3 might catalyze an uncharacterized homologue of the τm^5 2^U34 modification in mt-tRNAs. Interestingly, 5-carboxymethylaminomethyluridine (cmnm5U), was detected in mt-tRNAs in HeLa cells cultured under taurine-depleted conditions (Asano et al., 2018). cmnm5U is structurally similar to τm^5 2^U34 and occurs in the wobble position of a subset of bacterial tRNAs and in yeast and nematodes (Moukadiri et al., 2009). Bacterial MnmE and MnmG proteins (homologues of GTPBP3-MTO1) jointly use glycine (instead of taurine) as a substrate to synthesis cmnm5U modification. However, homologs of ELP3 are present in some bacteria and viruses, and their presence coincide with the lack of genes encoding for MnmE and MnmG in these organisms (Glatt et al., 2017) (McCloskey et al., 2001)(Moukadiri et al., 2009). Since τm^5 2^U34 is essentially synthesized when the taurine concentration is high, ELP3 likely localizes to mitochondria and catalyzes cmnm5U modification in mt-tRNAs at their U34 position under stress conditions (e.g., low concentrations of taurine).

Intriguingly, in yeast, ELP3-dependent tRNAs modification is necessary for mitochondrial function under stress conditions, and defects in this modification impaired mitochondrial protein synthesis, resulting in defective OXPHOS complexes (Tigano et al., 2015). While additional biochemical study is needed to fully understand the mechanistic role of mt-ELP3 in the formation of τm^5^U34 or its homologue, our data showed that the KAT domain of mt-ELP3 is essential for its mt-tRNAs function as evidenced by loss mt-tRNAs protection against angiogenin when introducing a mutation in its KAT domain.

The τm^5 2^U34 modification has a critical role in the efficiency and accuracy of mitochondrial translation. Indeed, loss of τm^5^U34 modification caused by MTO1 and/or a GTPBP3 defect resulted in a mitochondrial translation defect and contributed significantly to mitochondrial dysfunction (Ghezzi et al., 2012) (Kopajtich et al., 2014) (Asano et al., 2018). In the present study, an increase in the mitochondrial translation rate was observed in cells overexpressing mt-ELP3, consistent with our predicted function of mt-ELP3 in catalyzing the formation of τm^5^U34. Furthermore, the increased mitochondrial translation rate was associated with a substantial increase in the steady-state levels of OXPHOS complexes, resulting in an increase in mitochondrial ATP production.

Impaired canonical ELP3 tRNA modification activity is associated with several human diseases, including various cancers and neurodegenerative disorders (Hawer et al., 2019). For a full understanding of the role of mt-ELP3 in clinical manifestations of these diseases, studies of the tissue-specific expression of mt-ELP3 in human tissues are necessary. Our preliminary data indicate that mt-ELP3 is expressed in the brain at both the mRNA and protein levels (unpublished observations), but it is also expressed in several of the other tissues, although at lower levels. In agreement with these data, ELP3 was detected in the mitochondrial fraction of mouse brain (Stilger & Sullivan, 2013), suggesting that it is important in this tissue.

In brief, we present evidence that ELP3 localizes to mitochondria and functions as a non-canonical mt-tRNA modifying enzyme. This study provides a framework for further investigation that will clarify mechanistic insights into mt-ELP3 in the mt-tRNA modification reaction and the contributions of its defect in neurodegenerative diseases.

## Materials and Methods

### Cell culture

Human HEK-293T and HeLa cells were obtained from ATCC and grown in full medium: DMEM and EMEM, respectively, supplemented with 10% fetal bovine serum (FBS) (Gibco) and 1% penicillin/streptomycin (Gibco) at 37 °C and 5% CO_2_.

### Construction of plasmids and plasmid transfection

For the generation of pcDNA ELP3 and pcDNA mt-ELP3, the corresponding ORFs were amplified by PCR and cloned into a pCDNA4.T0 vector with a Flag tag at the N terminus.

The KAT domain mutant of ELP3 (pcDNA mt-ELP3 mut) was generated by PCR-based mutagenesis using specific primers harboring the desired mutation and pcDNA mt-ELP3 was used as a template. The mutation was confirmed by sequencing (Figure S4). The primers used for generation of this mutant were as follows: KAT-F, 5′-CAAACTGATGCTGAAATTTAGTAGG -3′; KAT-R, 5′-CTTTCATGCTGCTGATGGAGGAAGC-3′.

For GTPBP3 plasmid, GTPBP3 (NM_001128855) Human Tagged ORF Clone was purchased from Origene (cat # RC225798).

For the transfection assay, cells were transiently transfected with control siRNA duplex (OriGene #SR30002) and two Stealth siRNAs targeting ELP3 (ELP3 siRNA1(OriGene#SR310519A) and ELP3 siRNA 2 (OriGene#SR310519B)) or with pcDNA 4/T0 derivative plasmid expressing canonical ELP3 (pcDNA 4/T0-ELP3), mt-ELP3 (pcDNA 4/T0-mt-ELP3), mutant mt-ELP3 (pcDNA 4/T0-mt-ELP3) or empty pcDNA 4/T0 (pcDNA 4/T0) with Lipofectamine 3000 reagent, according to the manufacturer’s instructions. The cells were processed after 2 days of transfection.

### Generation of inducible HEK293 Cell Lines

Inducible HEK293 cell lines expressing mt-ELP3 were generated by using the Flp-In T-REx Core Kit from Invitrogen. Briefly, the cDNA encoding the mt-ELP3 was subcloned into the inducible expression vector, pcDNA5/FRT/TO using pcDNA mt-ELP3 as a template. Flp-In T-REx-293 cells were then co-transfected with the inducible expression vector pcDNA5/FRT/TO containing the mt-ELP3 cDNA and the pOG44 vector encoding the Flp recombinase by using OptiMEM/Lipofectamine2000 according to the manufacturer’s instructions. After 48h of transfection, cells were washed with PBS and incubated with selective medium containing 200 mg/ml hygromycin. The selective medium was changed at regular intervals until the desired number of cells was grown. Thereafter, the hygromycin-resistant cells were maintained in DMEM containing 10% FBS, 5ug/ml Blasticidin and 250ug/ml zeocin.

The expression of mt-ELP3 was induced by the addition of tetracycline (1ug/ml) to the culture medium for 24h.

### RNA isolation and APM-northern blotting analysis

Total RNA isolation was carried out using Trizol reagent (Invitrogen), according to the manufacturer’s recommendations. Northern blotting of APM-containing gels was performed as described to assess the thiolation status of tRNAs (Meseguer et al., 2015). Briefly, total RNA (5 µg) was run on a 15% polyacrylamide gel containing 7M urea and 10 µg/ml APM and then transferred to positively charged nylon membranes (Roche). Pre-hybridization and hybridization steps were performed with Dig Easy Hyb (Roche), according to the manufacturer’s instructions. The presence of tRNA species was detected with a specific DIG-labeled synthetic oligodeoxynucleotide probes **(Table S1).**

### In vitro angiogenin assay

Angiogenin nuclease-sensitivity assays were performed essentially as described (Rachid Boutoual et al., 2018). In brief, 1 µg of total RNA were mixed with 2.5 µg/ml recombinant angiogenin (ANG) in buffer (30 mM HEPES, pH 7.4, 30 mM NaCl, 0.01% BSA). Mixtures were incubated at 37°C for the indicated times. Angiogenin was inactivated by adding 5 µl of Gel Loading Buffer II (Life Technologies). The digested RNA samples were separated on 15% polyacrylamide, 8M urea gels and transferred to positively charged nylon membranes. The RNA was cross-linked to the membrane (60°C, 1 h) using freshly prepared EDC cross-linking solution (Kim et al., 2010)(Pall et al., 2007). Pre-hybridization and hybridization were performed with Dig Easy Hyb (Roche), according to the manufacturer’s instructions. tRNA species were detected with specific DIG-labeled synthetic oligodeoxynucleotide probes **(Table S1).**

### Blue native electrophoresis analysis

BN-PAGE was carried out as detailed (Sasarman et al., 2011). Samples containing 15 µg of protein were separated on 3–12% Bis-Tris Novex NativePAGE gel (Life Technologies). The relative levels of the assembled respiratory complexes I-V were assessed by western blot with following antibodies: anti-ND1(Thermo Fisher, 19703-1-AP), anti-SDHA (Cell signaling, 5839), anti- MTCO1(Thermo Fisher, 459600), anti-ATP5a (Abcam, 11448) and anti-Cytb (Abclonal, A17966). Horseradish-peroxidase-conjugated anti-rabbit IgG (Sigma, A6154) and anti-mouse IgG (Sigma, A4416) antibodies were used as secondary antibodies and protein signals were detected using the ECL system. Bis-Tris Novex NativePAGE (3–12%) gels were stained using Coomassie protein staining solution (50% methanol, 7% acetic acid and 0.1% Coomassie Brilliant Blue R stain).

### Western blots and subcellular fractionation

HEK293T cells were collected by centrifuging at 500xg for 5 min. Cells were lysed in RIPA buffer, and the homogenates were cleared by centrifugation and analyzed for protein concentration using BCA kit (Thermo Scientific). Protein lysates were separated on 4-20% gradient gels and transferred to PVDF membranes.

Subcellular fractionation was carried out as described with a slight modification (Park et al., 2013). Cells were gently homogenized in hypotonic buffer (10 mM Hepes, pH 7.9, 10 mM KCl, 0.1 mM EDTA) containing protease inhibitors (0.5 mM PMSF and protease inhibitor complete (Roche)) for 30 min on ice with the use of a dounce homogenizer until the vast majority of cells were ruptured and the nuclei were stained by Trypan blue, and then centrifuged at 800xg for 5 min at 4°C to obtain a pellet (nuclear fraction). The supernatant was centrifuged at 15,000xg for 10 min at 4°C to obtain a mitochondrial fraction (pellet) and cytosolic fraction (supernatant). The mitochondrial pellet was washed and suspended in hypotonic buffer. Protein concentration and immunoblotting were done as described above. Where indicated, the mitochondrial fraction was incubated with proteinase K (0.5–2 mg/ml) in hypotonic buffer in the presence of increasing concentrations of digitonin or proteinase K with 0.3% Triton X-100 for 15 min at 37 °C. After 15 min, the digestion was ended by adding 5 µL of 200 mM PMSF. The samples were boiled for 5 min and analyzed by western blotting.

For immunodetection, the following antibodies were used: anti-ELP3 (cell signaling, 5728S), anti- COX IV (cell signaling, 4850T), anti-GAPDH (Abcam, AB9484), anti-HSP60 (cell signaling, 4870S), anti-TOM20 (cell signaling, 42406), anti-Flag (Sigma, F1804-200UG), anti-acetylated- lysine (cell signaling, 9441) and anti-VDAC (cell signaling,4661).

### Immunofluorescence

HeLa cells were cultured on coverslips in 24-well plates overnight. After transient transfections, cells were washed with PBS, fixed with 4% paraformaldehyde–PBS for 20min at RT, washed with PBS, permeabilized with 0.3% Triton X-100 in PBS for 15 min and washed again with PBS. After being blocked with a solution containing 2% BSA, 0.05% Triton X-100 in PBS for 30 min at room temperature (RT), cells were incubated with anti-COX IV and anti-Flag antibodies in blocking solution for 1 h at RT. Upon washing with blocking solution, bound antibodies were subsequently detected by incubation, as appropriate, AlexaFluor 594-conjugated anti-rabbit (11012, Invitrogen) and AlexaFluor 488-conjugated anti-mouse (A11001, Invitrogen) secondary antibodies in blocking solution for 1h at RT. After staining with 1 mg/mL DAPI in PBS, cells were examined by using a Leica confocal microscope.

### Co-immunoprecipitation

293HEKT cells were co-transfected with indicated plasmids. After 72h post-transfection, transfected cells were harvested and lysed with IP lysis buffer (thermos scientific). The supernatants were obtained by centrifugation at 16,000 × *g* for 10 min at 4°C. Coimmunoprecipitation was performed using Dynabeads Protein G immunoprecipitation kit (Life Technologies) following the manufacturer’s instructions. Briefly, 10µg of anti-Flag M2 antibody was incubated with protein G Dynabeads for 10 minutes at RT. After washing, the antibody- Dynabeads complex was incubated (1 hour) with 300µg protein lysates (supernatant). Following washes, immune complexes were eluted in 20µl elution buffer and analyzed by immunoblotting.

Dynabeads were also incubated with anti-Flag M2 antibody (without lysates) and with lysates (without anti-Flag M2 antibody) and were used as negative control.

### De novo mitochondrial protein synthesis

De novo mitochondrial protein synthesis was performed as described (Zorkau et al., 2020). HEK239T cells were seeded in lab-tek chamber slides (ThermoFisher Scientific) and transfected with pcDNA-T0 or Flag-tagged pcDNA-mt-ELP3. After transfection, cells were incubated in methionine-free DMEM for 30 min. Then, the medium was replaced by methionine-free DMEM containing the methionine analogue L-homopropargylglycine (HPG), and cytosolic and mitochondrial translation were inhibited with 50 µg/ml cycloheximide (CHX) and 100 µg/ml chloramphenicol (CHL), respectively. After labelling period, slides were immediately placed on ice, and cells were pre-permeabilized with 0.005% digitonin in buffer A (10 mM HEPES/KOH, 10 mM NaCl, 5 mM MgCl_2_ and 300 mM sucrose in H_2_0, pH 7.5) for 2 min at RT before being fixed in pre-warmed 8% formaldehyde in buffer A for 15 min. Then, cells were fully permeabilized with 0.5% Triton X-100 in PBS. Pulse-labelled mitochondrial proteins were detected with a copper-catalyzed azide–alkyne cycloaddition (600 mM copper sulphate, 1.2 mM BTTAA, 40 µM picolyl Alexa Fluor 594 (CLK-1296-1) azide and 2 mM sodium ascorbate in PBS) by a click chemistry reaction for 1h at RT.

Samples were imaged using LSM700 confocal microscope. After imaging, each HPG-stained cell was selected by drawing an outline near the visible edge. The mean pixel intensity was then calculated for each bounded region representing the cell. Selection and analysis were performed with custom software written in Python.

For fluorescence intensity measured by flow cytometry, Brilliant Violet 421™ anti-DYKDDDDK (Flag-Tag) antibody (Cat# 637322, BioLegend) at 1:50 dilution was also added to the click reaction to sort transfected cells expressing mt-ELP3. Cells were then washed with 1XPBS and resuspended in 100µl of 1xPBS to acquire on Flow cytometer. Cells were acquired on FACS Aria II (BD Biosciences) with HPG signal collected in the Cherry filter while the Flag was collected in BV421 filter. Post-acquisition fcs files were analyzed using Flow jo v10 software (BD Biosciences). Cells were plotted as bimodal distribution of Cherry (HPG) Vs BV421 (Flag), MFI (mean fluorescence intensity) of HPG was quantified on Flag+ and Flag- population.

### Seahorse XF assays

Oxygen consumption rate (OCR) was assessed by a Seahorse XF96 Analyzer. Cells were seeded in Seahorse XF96 microplates at 4x10^4^ cells/well and incubated at 37°C in a CO_2_-free incubator for 1 h in prepared medium (1.8 mM CaCL_2_,139 mM NaCl, 20 mM HEPES, 1 mM NaHCO_3_, 25 mM glucose,1 mM pyruvate and 4 mM glutamine, pH 7.4) prior to analysis. Bioenergetic profiling was performed by monitoring oxygen consumption at basal levels, followed by the sequential injection of the following inhibitors: 2 µM oligomycin, 0.5 µM carbonyl cyanide-4-(trifluoromethoxy)-phenylhydrazone, 4 µM antimycin and 4 µM myxothiazol. All measurements were normalized against total number of cells in each well via imaging stained nuclei.

### Proteomics Analysis

#### 1) Protein digestion, acylation enrichment and desalting

Three independent biological experiments (with additional technical replicates) of HEK 293T cells control cells (-Tetra) and cells overexpressing mitochondrial ELP3 (+Tetra) were investigated by acetyl enrichment followed by mass spectrometric analysis. Cell lysates were immersed in lysis buffer containing 8 M urea, 200 mM triethylammonium bicarbonate (TEAB), pH 8.5, 75 mM sodium chloride, 1 µM trichostatin A, 3 mM nicotinamide, and 1x protease/phosphatase inhibitor cocktail (Thermo Fisher Scientific, Waltham, MA), and homogenized for 2 cycles with a Bead Beater TissueLyser II (Qiagen, Germantown, MD) at 24 Hz for 3 min each. Lysates were clarified by spinning at 15,700 x *g* for 15 min at 4°C, and the supernatant containing the soluble proteins was collected. Protein concentrations were determined using a bicinchoninic acid protein (BCA) Assay (Thermo Fisher Scientific, Waltham, MA), and subsequently 1-2 mg of protein from each sample were brought to an equal volume using a solution of 8 M urea in 50 mM TEAB, pH 8. Proteins were reduced using 20 mM dithiothreitol (DTT) in 50 mM TEAB for 30 min at 37 °C, and after cooling to room temperature, alkylated with 40 mM iodoacetamide (IAA) in 50 mM TEAB for 30 min at room temperature in the dark. Samples were diluted 4-fold with 50 mM TEAB, pH 7.5, and proteins were digested overnight with a solution of sequencing-grade trypsin (Promega, San Luis Obispo, CA) in 50 mM TEAB at a 1:50 (wt:wt) enzyme:protein ratio at 37°C. This reaction was quenched with 1% formic acid (FA) and the sample was clarified by centrifugation at 2,000 x g for 10 min at room temperature. Clarified peptide samples were desalted with Oasis 10-mg Sorbent Cartridges (Waters, Milford, MA), and subsequently all desalted samples were vacuum dried. The digestions were re-suspended in 1.4 mL of immunoaffinity purification (IAP) buffer (Cell Signaling Technology, Danvers, MA) containing 50 mM 4- morpholinepropanesulfonic acid (MOPS)/sodium hydroxide, pH 7.2, 10 mM disodium phosphate, and 50 mM sodium chloride for PTM enrichment. Peptides were enriched for acetylation with anti-acetyl antibody conjugated to agarose beads (Acetyl-Lysine Motif Kit; Cell Signaling Technology, Danvers, MA). This process was performed according to the manufacturer protocol, and each sample was incubated overnight with half a vial of washed beads. Finally, acetyl-enriched peptides were eluted from the antibody-bead conjugates with 0.1% trifluoroacetic acid in water and were desalted using C-18 zip-tips (Millipore, Billerica, MA). Samples were vacuum dried and re-suspended in 0.2% FA in water.

#### 2) Mass Spectrometric Analysis

Samples were analyzed by reverse-phase HPLC-ESI-MS/MS using an Eksigent Ultra Plus nano- LC 2D HPLC system (Dublin, CA) with a cHiPLC system (Eksigent) which was directly connected to a quadrupole time-of-flight (QqTOF) TripleTOF 6600 (or TripleTOF 5600) mass spectrometer (SCIEX, Concord, CAN). After injection, peptide mixtures were loaded onto a C18 pre-column chip (200 µm x 0.4 mm ChromXP C18-CL chip, 3 µm, 120 Å, SCIEX) and washed at 1-2 µl/min for 10 min with the loading solvent (H_2_O/0.1% formic acid) for desalting. Subsequently, peptides were transferred to the 75 µm x 15 cm ChromXP C18-CL chip, 3 µm, 120 Å, (SCIEX), and eluted at a flow rate of 300 nL/min with 2-3 hour gradients using aqueous (A) and acetonitrile (B) solvent buffers.

**3) Data-dependent acquisition (DDA)** to identify acetylated peptides and build spectral libraries. Every cycle consisted of one 250 ms precursor ion scan followed by isolating the top 30 most abundant precursor ions between 400-1,500 m/z (tandem mass spectra accumulation time of 100 ms yielding a total cycle time of 3.25 sec; ‘high sensitivity’ product ion scan mode, software: Analyst 1.7; build 96) as previously described (Schilling et al., 2015).

**4) Data-independent acquisition (DIA)** to quantify the PTM enriched peptides using the TripleTOF 6600 mass spectrometer. Every cycle consisted of one 250 ms precursor ion scan with 400-1,250 m/z mass range. Subsequently, windows of variable width are passed in incremental steps over the full mass range (m/z 400-1,250). The cycle time of 3.2 sec includes the 250 msec precursor ion scan followed by 45 msec accumulation time for each of the variable 64 DIA segments (variable windows isolation scheme (Schilling et al., 2017)) monitoring fragment ion masses between 100-1,500 m/z.

**5) Mass-spectrometric data processing and quantification.** Mass spectrometric data-dependent acquisitions (DDA) were analyzed using the database search engine ProteinPilot (SCIEX Beta 4.5, revision 1656) using the Paragon algorithm (4.5.0.0,1654). Using these database search engine results MS/MS spectral libraries were generated. Quantification was performed in Skyline using MS1 Filtering and DIA/SWATH MS2 data quantification (using XICs of 6-10 MS/MS fragment ions, typically y- and b-ions, matching to specific peptides present in the spectral libraries) as previously described (Schilling et al., 2012)(Rardin et al., 2015)(Gut et al., 2020).

**6) Data availability.** The mass spectrometric raw data are deposited at ftp://massive.ucsd.edu with the MassIVE ID MSV000088668 (password winter): Go to http://massive.ucsd.edu/ProteoSAFe/status.jsp?task=3b6b233db1d44e65aee8956d17eb7289 Enter the username and password in the upper right corner of the page:

Username: MSV000088668_reviewer; Password: winter

The raw data is also available at ProteomeX change with the ID PXD030850 (will be public upon release).

### Statistical analysis

The statistical analyses were performed using Graph Pad Prism 9. Student’s t-test was used in all comparisons of data. The statistically significant differences between the means were indicated by asterisks (*p < 0.05, **p < 0.01 or ***p < 0.001), and non-significant differences by n.s.

## Acknowledgements

This project was supported by Buck Institute for Research on Aging. We acknowledge the support of instrumentation for the TripleTOF 6600 system from the NCRR shared instrumentation grant 1S10 OD016281 (Buck Institute). We thank Gary Howard for editing the manuscript and John CW Carroll for his help with the figures and graphics.

---

**Figure S1.**
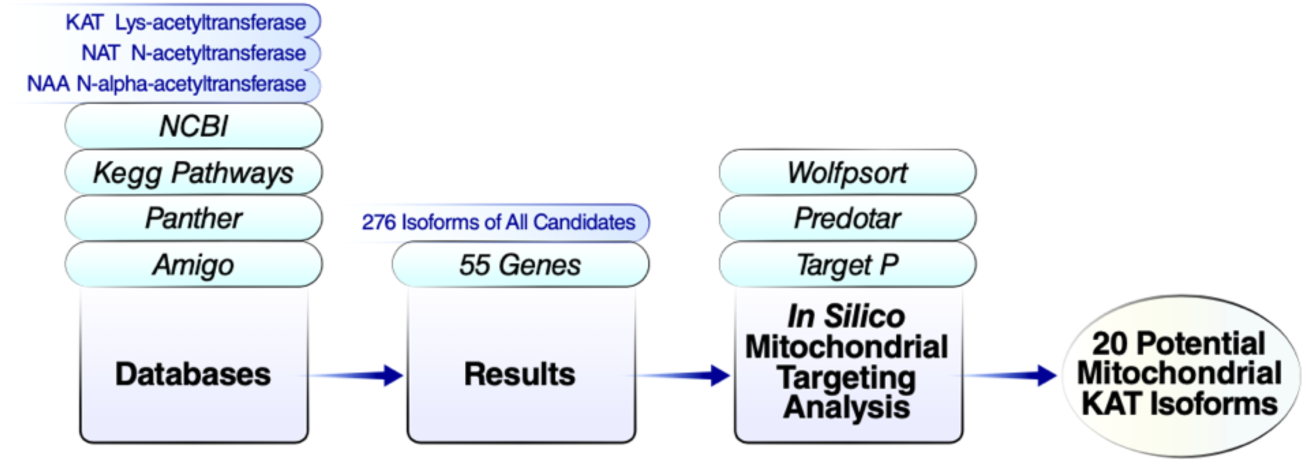
The methodology used to identify putative mitochondrial acetyltransfereases.

**Figure S2.**
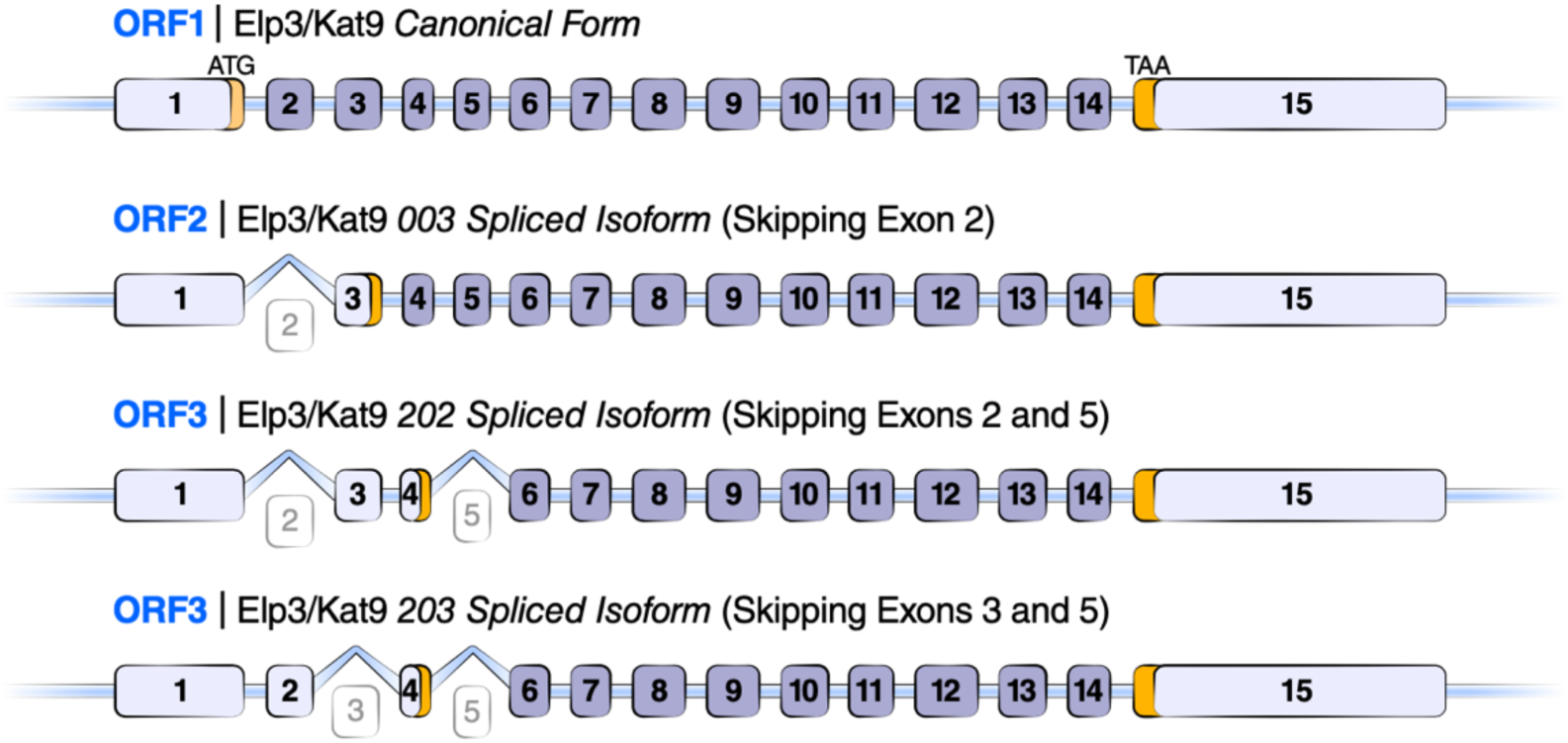
Representation of full-length ELP3 and its spliced isoforms. Start and stop codon are highlighted in each isoform.

**Figure S3.**
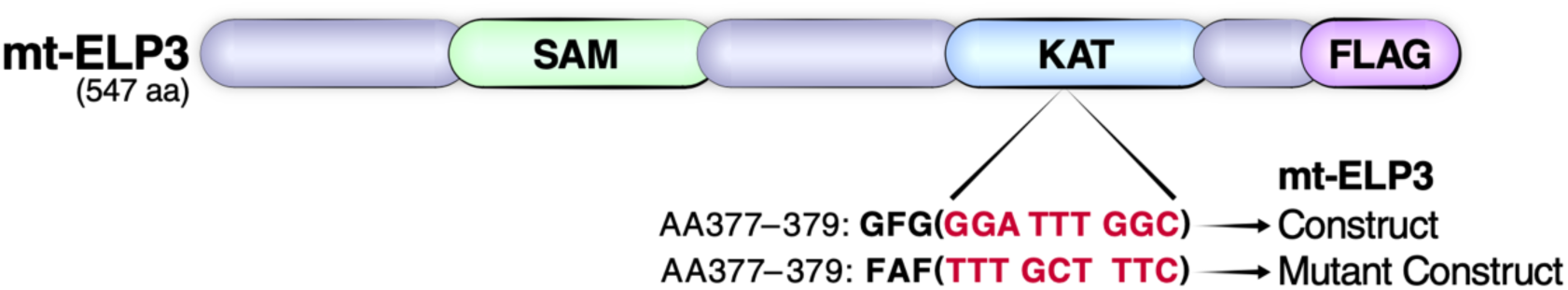
Schematic view of the domain structure of mt-ELP3 construct. The position of the change introduced in mt-ELP3 to generate mt-ELP3 mutant is highlighted.

**Figure S4.**
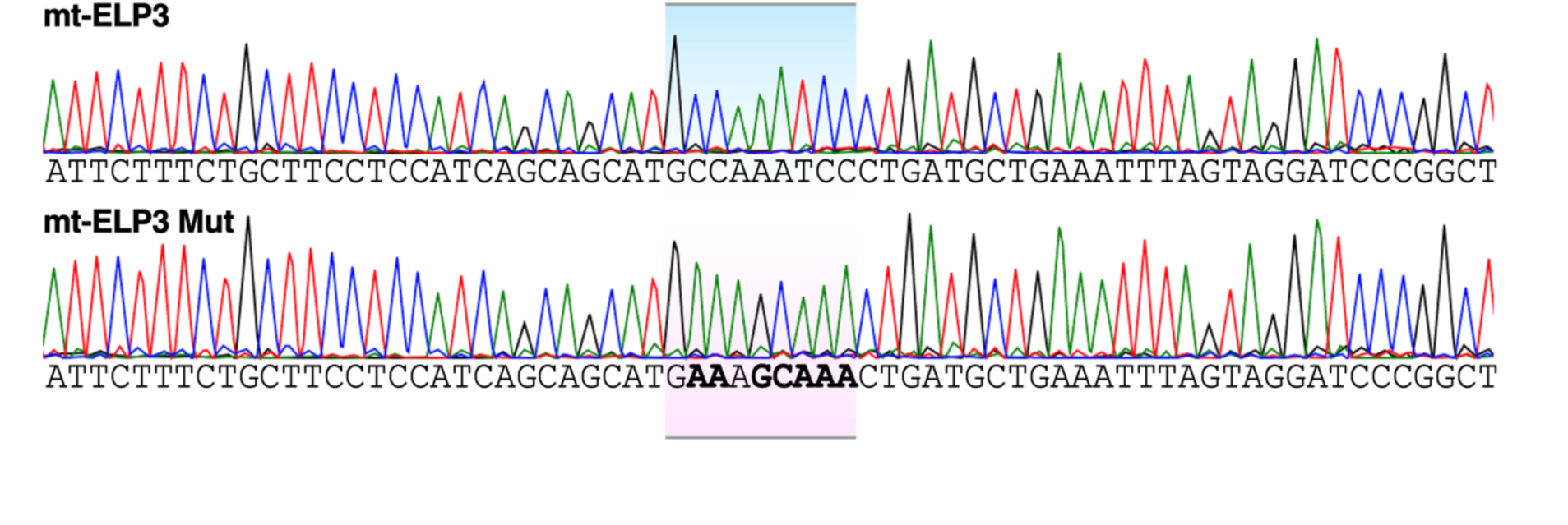
Part of the sequencing results. The mutation: amino acid (AA) 377-379: GFG(GGATTTGGC) è FAF(TTTGCTTTC) is highlighted.

**Figure S5.**
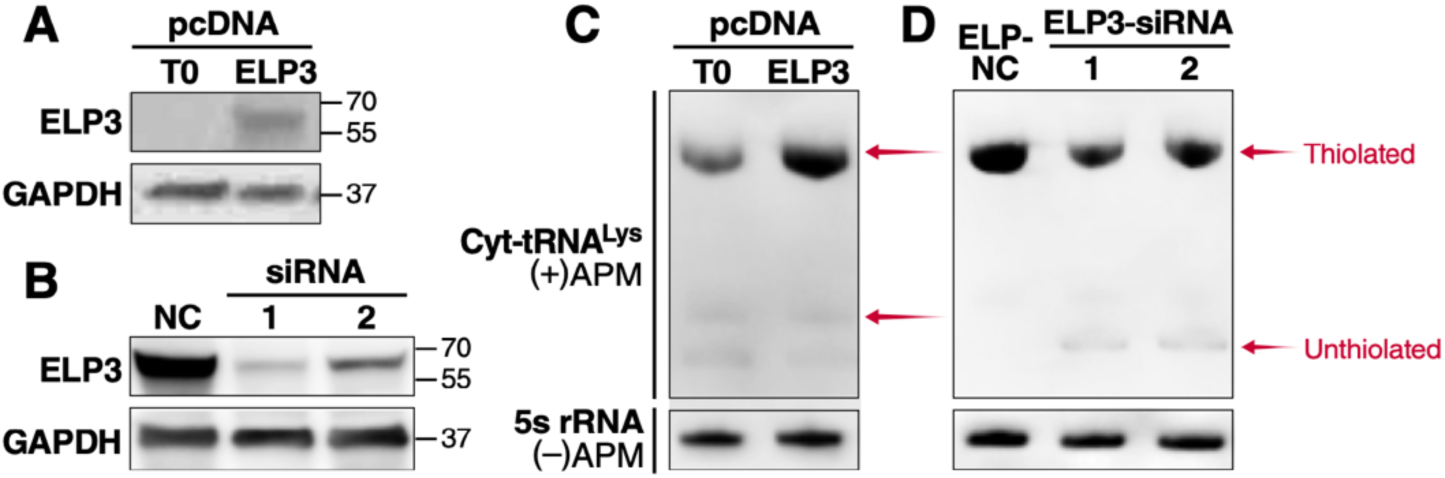
Overexpression or transient silencing of canonical ELP3 in HEK293T cells affects the 2 thiolation modification status of cyt-tRNA^Lys^ molecules. (**A and B**) Western blot analysis of canonical ELP3 expression in HEK293T cells transfected with empty plasmid (pcDNA4 T0) and pcDNA ELP3 **(A)** and in ELP3 siRNA 1, ELP3 siRNA 2- and Negative Control (NC) siRNA-transfected HEK293T cells **(B)**. The membranes were also probed with an antibody against GAPDH, which was used as a loading control. **(C and D)** APM-northern analysis of the 2-thiolation status of mt-tRNA^Lys^ molecules isolated from HEK293T transfected with pcDNA4 T0 and pcDNA ELP3 **(C)** and from ELP3 siRNA 1, ELP3 siRNA 2- and Negative Control (NC) siRNA-transfected HEK293T cells **(D)**. The same amount of total RNA (5 µg) was run in a denaturing polyacrylamide-urea gel in the presence (+) or absence (-) of APM. The thiolated tRNAs were detected as retarded bands in the presence of APM. The APM (−) membrane was probed with 5S rRNA as a loading control.

**Table S1.**
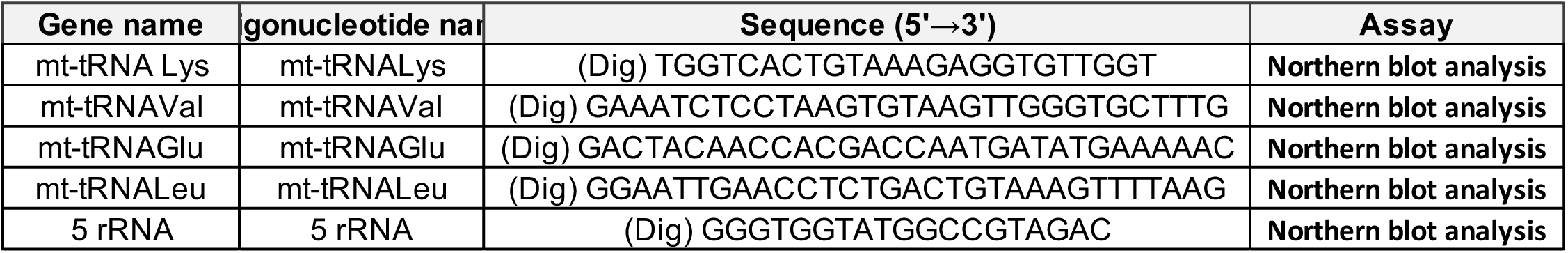
Digoxigenin (DIG)-labeled oligodeoxynucleotide probes

